# Generalizable predictive modeling of semantic processing ability from functional brain connectivity

**DOI:** 10.1101/2021.05.11.443600

**Authors:** Danting Meng, Suiping Wang, Patrick C.M. Wong, Gangyi Feng

**Author notes:** Corresponding author Gangyi Feng, Ph.D., Suiping Wang, Ph.D.

## Abstract

Semantic processing (SP) is one of the critical abilities of humans for representing and manipulating meaningful and conceptual information. Neuroimaging studies of SP typically collapse data from many subjects, but both its neural organization and behavioral performance vary between individuals. It is not yet understood whether and how the individual variabilities in neural organizations contribute to the individual differences in SP behaviors. Here we aim to identify the neural signatures underlying SP variabilities by analyzing individual functional connectivity (FC) patterns based on a large-sample Human Connectome Project (HCP) dataset and rigorous predictive modeling. We used a two-stage predictive modeling approach to build an internally cross-validated model and to test the model’s generalizability with unseen data from different HCP sub-populations and task states as well as other out-of-sample datasets that are independent of the HCP. FC patterns within a putative semantic brain network were significantly predictive of individual SP scores summarized from five semantic tasks. This cross-validated predictive model can be used to predict unseen HCP data. The model generalizability was enhanced with FCs in language tasks than resting state and other task states and was better for females than males. The model constructed from the HCP dataset can be generalized to two independent cohorts that participated in different semantic tasks. FCs connecting to the Perisylvian language network show the most reliable contributions to predictive modeling and the out-of-sample generalization. These findings contribute to our understanding of the neural sources of individual differences in SP, which potentially lay the foundation for personalized education and improve intervention practice for patients with SP and language deficits.

## Introduction

Making sense of the outside world is critical for human survival and development. The ability to store, retrieve, and manipulate meaningful information (e.g., concepts) is central to many cognitive functions and is also a defining characteristic of human brains (Berwick *et al*., 2013; Nation & Snowling, 1998; Ratcliff *et al*., 2010). Although this so-called semantic processing (SP) ability is a gift for humans in general, there are extensive inter-individual differences evidenced by various behavioral tasks, ranging from object recognition (Lewellen *et al*., 1993), conceptual categorization, lexical decision (Plaut & Booth, 2000; Schilling *et al*., 1998; Yap *et al*., 2012), to language comprehension and production (Just & Carpenter, 1987; Walker *et al*., 1994). These behavioral measures in SP have been also demonstrated to be associated with executive control (Allen *et al*., 2012), attention (McGlinchey-Berroth *et al*., 1993), selection (Nation & Snowling, 1998), and inhibition abilities (Cain, 2006), suggesting SP comprises multiple cognitive components where inter-individual variabilities in these functions may contribute to the variability in SP. Although these behavioral findings provide insights into our understanding of the composition of the SP variability, it is not yet clear to which extent the individual differences in neural organizations contribute to the individual differences in SP behaviors.

One candidate neural source underlying SP behavioral variabilities are inter-regional functional connectivity patterns within a putative semantic network (Binder *et al*., 2009; Binder & Fernandino, 2015). The multifaceted essence of SP in cognitive composition suggests that the neural implementation of SP requires joint efforts of multiple brain regions, not only reflecting in regional activities but also more in inter-regional connectivity. In consistence of this hypothesis, findings derived from a large number of group-level neuroimaging studies have shown that SP is robustly related to a distributed brain regions, including the left inferior frontal gyrus, left superior temporal gyrus, left middle temporal gyrus, left anterior temporal lobe, left angular gyrus, left inferior parietal lobule, medial prefrontal cortex, and posterior cingulate cortex (Binder & Desai, 2011; Binder *et al*., 2009). These SP-related regions can be divided into three sub-networks, classical Perisylvian language network (PSN), frontoparietal network (FPN), and default mode network (DMN) based on their functional connectivity profiles (Xu *et al*., 2016). The functional associations of these subnetworks are distinct and relate to different aspects of SP; for example, PSN regions have been proposed to be related to high-level linguistic processes (Fedorenko, 2014; Fedorenko *et al*., 2011; Feng *et al*., 2016), language learning (Forkel *et al*., 2014; Xiang *et al*., 2012), and language recovery after stroke (Dehaene-Lambertz *et al*., 2006; Griffis *et al*., 2017; López-Barroso *et al*., 2013; Ojemann, 1991; Saur *et al*., 2006); FPN regions relate to semantic control processes (Feng *et al*., 2016; Geranmayeh *et al*., 2012; Geranmayeh *et al*., 2017; Geranmayeh *et al*., 2014; Wirth *et al*., 2011), while DMN regions relate to social conceptual processing (Binder *et al*., 2009; Binder & Fernandino, 2015).

While this large body of research paints a convincing picture of the relationship between the semantic network and SP at the group level, it does not adequately acknowledge the tremendous individual differences in SP and few studies investigate whether participants’ variabilities in neural patterns contribute to predicting individual differences in SP behaviors, and if so, how. With a small sample size and correlational approaches, previous studies showed that inter-individual variability of functional connectivity within the semantic network was associated with individual differences in SP (Turken & Dronkers, 2011; Xu *et al*., 2016). In particular, the strength of the connectivity between a network hub in the middle temporal gyrus and other regions explains inter-individual differences in SP performance (Krieger-Redwood *et al*., 2016; Mollo *et al*., 2016; Vatansever *et al*., 2017; Wei *et al*., 2012). These studies provide initial evidence supporting that neural connectivity acts as one major source of individual differences in SP, while there are significant limitations of the findings due to the methodology constraint and solely focusing on individual connectivity strengths.

Small sample size and traditional correlational approaches could potentially inflate the neural-behavioral correlations, which limits the generalization of a finding derived from one population to other unseen samples. Questions were raised on the reliability and replicability of the correlational findings. Most of the previous neuroimaging studies in SP used small samples, which has low statistical power in general, leading to inflated estimates of effect size and poor replicability (Dubois & Adolphs, 2016; Schönbrodt & Perugini, 2013). For example, for behavioral studies, statistical simulation with the Monte-Carlo procedure has demonstrated that stable correlation estimation should approach a minimal sample size of 250 for a significant degree of confidence (Schönbrodt & Perugini, 2013). For fMRI studies, small sample sizes reduce the stability, test-retest reliability, and replicability of an activation effect across paradigms used (Bossier *et al*., 2020; Kühberger *et al*., 2014). In addition to the effect of sample size, traditional correlational methods often overestimate the relationships and do not ensure the generalizability of the established relationship from a sample population to out-of-sample subjects (Dubois & Adolphs, 2016; Lo *et al*., 2015; Whelan & Garavan, 2014). This out-of-sample generalization ability is rarely demonstrated in both behavioral and neuroimaging studies, which may partly be due to the small sample size and lack of using machine-learning approaches. For example, previous studies demonstrated that machine learning algorithms, cross-validation, and randomization procedure can prevent overfitting data and promote model generalization and replication (Shmueli, 2010; Yarkoni & Westfall, 2017).

To identify the functional connectivity patterns underlying individual differences in SP while overcoming the above-mentioned methodological limitations, here we used the state-of-art predictive modeling approach (Finn *et al*., 2015; M. Rosenberg *et al*., 2013; Shen *et al*., 2017) with two-step cross-validation and generalization procedures to construct and validate SP predictive models based on a large number of subjects (*N* = 868) from the Human Connectome Project (Van Essen *et al*., 2013). First, to estimate the individual differences in SP while minimizing the bias in selecting a single semantic task, we extracted a latent SP component from five behavioral tasks related to semantic processing with confirmation factor analysis. Secondly, we constructed cross-validated semantic models to predict the latent SP scores with half of the HCP dataset. We identified the most predictive functional connections to build a predictive model, and the model was then used to generalize to another half of the HCP samples as well as to two other independent datasets with different populations and tasks. The overall model prediction and generalizability were evaluated by bootstrapping and permutation procedures. Moreover, previous studies have identified factors that modulate the predictability of individual behavioral scores and cognitive traits with functional connectivity patterns (Greene *et al*., 2018; R. Jiang *et al*., 2020). Here, we further explored to which extent the populational factors (i.e., gender and age) and task status during fMRI imaging modulate the model prediction and generalization.

## Materials and Methods

### Dataset description and sample population

Three datasets were used to construct and cross-validate predictive models and assess model generalization performance. The three datasets include the Human Connectome Project (HCP) 1200 Subjects Release (Van Essen *et al*., 2013), the semantic lexical decision (SLD) dataset (Feng *et al*., 2016), and the Alice story comprehension (ASC) dataset (Bhattasali *et al*., 2020). The HCP dataset was used to construct and cross-validate models and assess in-sample model generalization. Another two independent datasets were used to estimate the out-of-sample generalization performance of the HCP models. The details of the three datasets are described below.

#### HCP dataset

The HCP dataset includes behavioral and 3T MR imaging data from 1206 healthy young adult subjects. These subjects were asked to complete a resting-state session and a range of cognitive tasks during fMRI scanning (Barch *et al*., 2013). They also completed various cognitive assessments outside the scanner. Detailed descriptions of these behavioral and fMRI tasks were listed in Supplementary Methods and Table S1 (*Supplementary Information, SI*). We applied the following criteria to exclude subjects from the functional connectivity (FC) analysis and predictive modeling: *i*) Subjects must have completed all resting-state and task-based fMRI scanning (including working memory, gambling, motor, language, social cognition, relation, and emotion processing tasks) as well as cognitive assessments of interest (i.e., picture vocabulary test, oral reading recognition test, pattern completing processing speed test, picture sequence memory test, list sorting test, flanker tasks test, dimensional change card sort test, 2-minute walk test, 4-meter walk test, 9-hole pegboard test, and grip strength dynamometry test; see Table S1 for detailed description). Subjects with any missing data in any of the scans and tests were discarded (*N* = 253); *ii*) Imaging quality control. We excluded subjects whose data collected during the period of known intermittent problems with head coil leading to temporal instability in acquisitions (i.e., QC_Issue = C; *N* = 75); *iii*) We also removed subjects whose one or more functional scans contained any significant coil- or movement-related artifact that manifests prominently in the “minimally preprocessed” data (i.e., QC_Issue = D; *N* = 10). Finally, 868 subjects (407 males; age range = 22–35 years old) were included in our analysis. The HCP scan protocol was approved by the local Institutional Review Board at Washington University in St. Louis.

#### Independent datasets

Two independent datasets (i.e., SLD and ASC datasets) were included to evaluate out-of-sample model generalization. These two datasets were chosen because one uses classical semantic processing manipulation (semantic priming) with rigorous experimental controls while another uses a more natural setting probing language comprehension. The SLD dataset has both resting-state and task-based fMRI scans. This dataset consists of 26 Chinese healthy subjects (11 males, 18–28 years old) (Feng *et al*., 2016). All subjects were right-handed, with normal or corrected-to-normal vision, and no prior history of neuropsychiatric disorders. During the resting-state scans, subjects were asked to fixate on a cross and lie still. For the task-based scanning, the subjects were asked to complete a lexical decision task for a list of Chinese word pairs (i.e., judging the second word is a real word or not). Word pairs include semantic-related (e.g., “Bread – Cake”), unrelated (e.g., “Driver – Cake”), and nonword pairs. The ASC dataset only includes task-based fMRI scans (Bhattasali *et al*., 2020). This dataset includes 26 healthy native English speakers (11 males, 18–24 years old, right-handed). During the scanning, participants listened passively to an audio storybook reading by Kristen McQuillan’s first chapter of *Alice’s Adventure in Wonderland* (duration = 12.4 min). Participants completed a twelve-question multiple-choice questionnaire concerning events and situations described in the story after scanning.

### Imaging data acquisition

*HCP dataset*. Structural T1-weighted images were acquired using a magnetization-prepared rapid acquisition gradient-echo (MPRAGE) sequence (TR = 2400 ms, TE = 2.14 ms, flip angle = 8^°^, FOV = 228 × 224 × 180 mm, 0.7-mm isotropic voxels). Whole-brain gradient echo-planar imaging (EPI) acquisitions were acquired with a 32 channel head coil on a modified 3 T Siemens Skyra with TR = 720 ms, TE = 33.1 ms, flip angle = 52^°^, FOV = 208 × 180 × 144 mm, 2.0-mm isotropic voxels, and 72 oblique axial slices that alternated between phase encoding in a right to left direction in one run and phase encoding in a left to the right direction in the other run, with a multi-band acceleration factor of 8 (Uğurbil *et al*., 2013; Van Essen *et al*., 2012). Imaging data with two-phase encoding directions were used to calculate functional connectivity. One session of resting-state scan and seven sessions of the task-state scans were acquired separately (see more details in Supplementary Methods).

#### SLD dataset

This data was acquired using a Siemens Trio 3T MRI system with a 32-channel head coil. T1-weighted high-resolution structural images were acquired using a MPRAGE sequence (176 slices, TR = 1900 ms, TE = 2.53 ms, flip angle = 9^°^, voxel size = 1 × 1 × 1 mm). The functional data were recorded by a T2*-weighted EPI pulse sequence (38 slices, TR = 2000 ms, TE = 20 ms, flip angle = 90^°^, field of view = 224 mm × 224 mm, in-plane resolution = 3.5 × 3.5 mm, slice thickness = 3.5 mm with 1.1 mm gap). A total of 240 volumes were collected for the resting-state scans and 270 volumes for the task-state scans.

#### ASC dataset

The neuroimaging data were acquired using a 3T MRI scanner (Discovery MR750, GE Healthcare, Milwaukee, WI) with a 32-channel head coil. Anatomical images were collected with a high-resolution T1-weighted with an MPRAGE sequence (voxel size = 1 × 1 × 1 mm). The functional data for ten subjects were recorded by the T2∗-weighted EPI sequence (44 slices, TR = 2000 ms, TE = 27 ms, flip angle = 77°, field of view = 216 × 216 mm, voxel size = 3 × 3× 3 mm, acceleration factor = 2). Sixteen subjects were scanned with a three-echo EPI sequence (33 slices, TR = 2000 ms, TE = 27.5 ms, field of view = 240 mm × 240 mm, in-plane resolution = 3.75 × 3.75, slice thickness = 3.8 mm with 0.05 mm gap).

### Estimation of latent semantic processing (SP) ability

#### HCP dataset

Five behavioral assessments were selected to estimate a latent SP. These behavioral tests were chosen from the HCP cognitive battery tested by the NIH Toolbox Cognition Battery (Gershon et al., 2013). We select tasks that require participants to store, retrieve, and/or manipulate meaningful object or conceptual information. The five tasks include the oral reading recognition test ([ReadEng], measuring the language processing and reading decoding ability), the picture vocabulary test ([PicVoc], measuring the language/vocabulary comprehension ability), pattern comparison processing speed ([ProcSpeed], measuring processing speed of pictures discernment), picture sequence memory ([PicSeq], measuring episodic memory ability of illustrated objects and activities), and list sorting ([ListSort], measuring temporal storage performance of sequencing different visually- and orally-presented objects).

We also included two tasks related to general cognitive control (CC) and four tasks related to motor control (CC) to estimate the latent components related to CC and MC, respectively. The two CC-related tests include dimensional change card sort ([CardSort], measuring executive function and cognitive flexibility) and flanker task ([Flanker], measuring selection and inhibition ability). The four MC-related tests include the 2-minute walk test ([Endurance], measuring endurance), 4-meter walk test ([GaitSpeed], measuring locomotion), 9-hole pegboard ([Dexterity], measuring dexterity), and grip strength dynamometry ([Strength], measuring strength). These tests do not include any meaningful object or conceptual stimuli.

To estimate latent variables related to SP, CC, and MC ability individually, we adopted a 3-factor confirmatory factor analysis (CFA) with those test variables for all selected subjects. The CFA is a multivariate statistical procedure which often used to test how well the measured variables contribute to the latent constructs of interest. We conducted the CFA model with R package Lavaan (Rosseel, 2012).

#### SLD and ASC datasets

For the SLD dataset, we used the semantic priming effect in lexical decision time as individuals’ SP scores. The strength of semantic priming was calculated by subtracting the lexical decision time of the unrelated condition (a target word paired with an unrelated prime) from the semantic-related condition (a target word paired with a semantically related condition) for each subject. Higher priming scores reflect more facilitation in semantic processes (e.g., semantic access) of the target words due to the presence of semantically related primes. Trials with error responses and response times deviating from the mean of all trials by more than three standard deviations were removed. The semantic priming scores across subjects range from − 66.68 ms to 87 ms with a mean of 21.02 ms. For the ASC dataset, the performance of the story comprehension test was used as individuals’ SP scores. These comprehension scores range from 5 to 12 with a mean of 9.81 (*SD* = 1.67).

### Imaging data analysis

#### Preprocessing

All three datasets were processed with the same analysis pipeline. The HCP data was downloaded in its minimally preprocessed form (i.e., after motion correction, B0 distortion correction, coregistration to T1-weighted images, and normalization to MNI152 space; see Glasser *et al*. (2013) for detailed preprocessing parameters). The other two datasets were preprocessed with the same procedure as the HCP dataset with Data Processing Assistant for Resting-State fMRI (DPARSF) (Yan & Zang, 2010). Before calculating functional connectivity matrices, the preprocessed datasets were resliced into 3.5 mm^3^ voxel size. We applied band-pass filtering (0.01–0.08 Hz) to remove physiological noises and linear detrend to remove slow drifts. We then regressed out variances of nuisance variables including 12 head motion parameters, global signals, white-matter signals averaged from the deep cerebral white matter, cerebrospinal fluid signals averaged from the ventricles to reduce non-neuronal contributions to variable covariance.

### Functional connectivity matrix construction

To derive functional connectivity patterns within the semantic network for each subject, we used a semantic brain template of 60 nodes. This template is constructed based on a meta-analysis deriving from 120 SP-related fMRI studies from Binder *et al*. (2009). This brain template includes the frequently activated areas summarized from a range of semantic manipulations (e.g., words versus pseudowords, semantic versus phonological tasks, and high versus low meaningfulness conditions, etc.). All the regions are listed in Table S2 (SI). For each region, we defined a sphere with a radius of 6 mm. Representative mean time series for each region were extracted by averaging all voxels within that region. The inter-regional functional connectivity was measured by calculating the pairwise Pearson correlation coefficient of the time series between each pair of regions. The functional connectivity was then normalized with Fisher’s *r*-to-*z* transformation. Finally, for each subject, a 60×60 symmetric connectivity matrix was obtained for each scanning session. For the HCP dataset, each subject had eight connectivity matrixes, separately calculated from the resting-state, working memory, gambling, motor, language, social cognition, relation processing, and emotion processing task fMRI. For the SLD dataset, each subject has two connectivity matrixes (i.e., the resting-state and SLD task). For the ASC dataset, each subject has only one connectivity matrix (i.e., the story comprehension task).

### Predictive model construction procedure (Phase 1)

We randomly split the HCP sample into two parts (*N* = 434 each) for predictive model construction and in-sample generalization estimation, respectively (see the overview of the analysis schema in Fig. 1). To construct a cross-validated SP prediction model, we combined the 10-fold cross-validation with bootstrapping and permutation procedures (Feng *et al*., 2018; Feng *et al*., 2021). First, we randomly split the subjects (*N* = 434) into 10 folds. Nine folds of the subjects (i.e., training set) were used for model training and the held-out fold (i.e., testing set) was used for model validation. To avoid overfitting the model with a large number of edges, we employed a feature selection procedure to select the most informative features during modeling training. For each training set, we computed the partial Pearson correlations between functional connectivity strengths and SP scores for each edge, where age and gender were controlled for. We discarded the non-significant edges with a threshold of *P* = 0.005 while selected the positively and negatively correlated edges separately, as predictive features to train models. The two types of edges were modeled separately because their connection patterns could contribute differently to the model prediction based on previous findings (Feng *et al*., 2021; M. D. Rosenberg *et al*., 2016). A different feature selection threshold of 0.001 or 1% of the total edges was also used to ensure the reliability of the model prediction. Next, to further reduce the number of feature dimensions, we used principal component analysis (PCA) to summarize the main components and select the components that were significantly correlated with the SP scores with a threshold of *P* = 0.05. These feature selection and dimension reduction procedures were conducted only in each training set that is independent of the held-out testing set. Finally, a linear regression model was constructed with the selected principal components as predictive features and the SP score as the dependent variable. This trained model was then applied to the held-out 10% unseen subjects to predict their SP scores. This model validation process was repeated 10 times so that all the subjects’ SP scores were predicted. Pearson correlations between the observed and predicted SP scores were used to assess the model prediction performance (i.e., *r*_[observed, predicted]_).

**Fig. 1.**
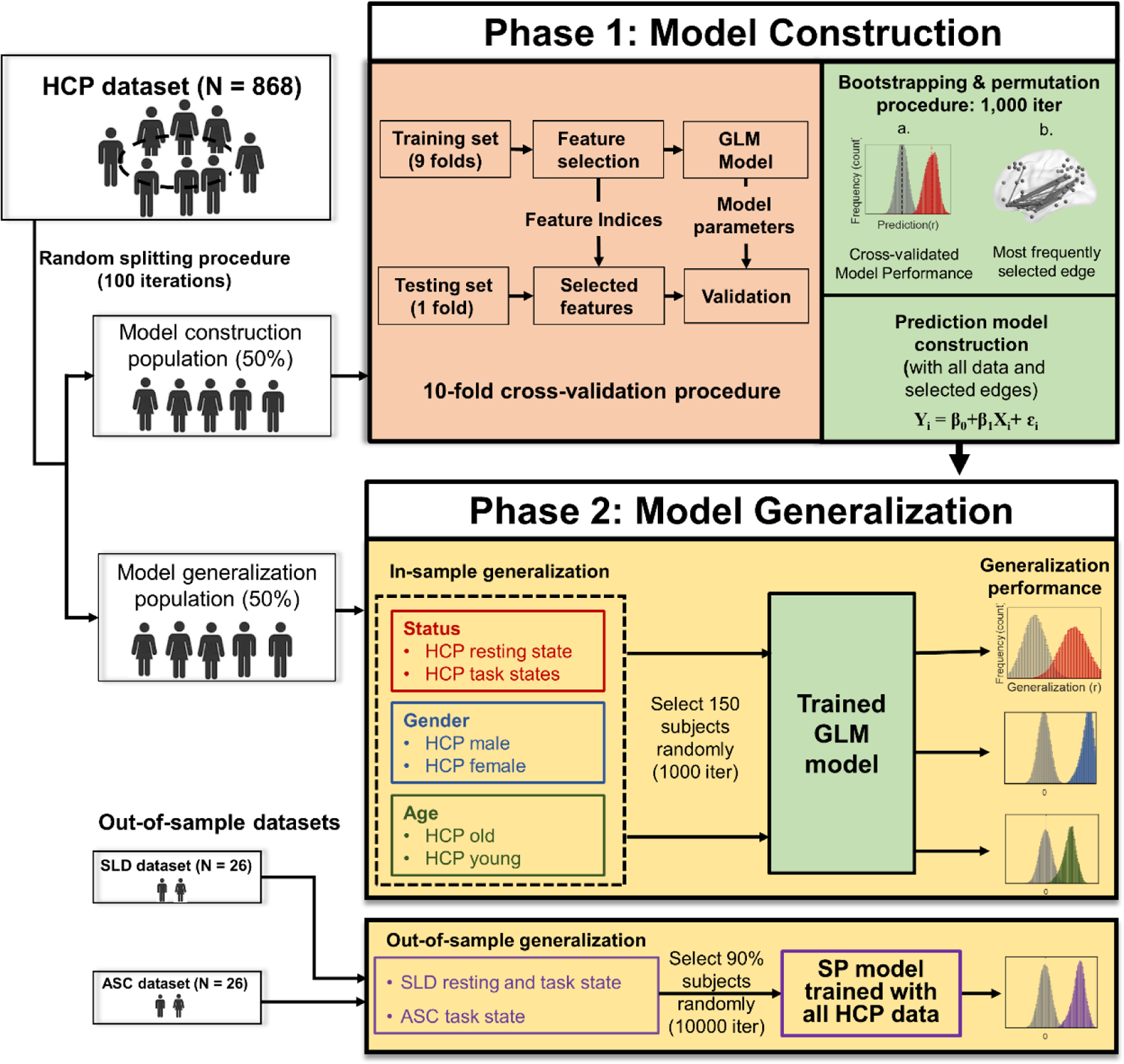
Schema of the predictive modeling and generalization evaluation procedure. The HCP subjects were randomly split into two groups (50% each) for model construction (Phase 1) and generalization evaluation (Phase 2), respectively. This random splitting procedure was repeated 100 times. For each splitting, at the model construction phase, we used 10-fold cross-validation (light red box), bootstrapping, and permutation procedures to construct and validate models (green box). At the end of Phase 1, a prediction model was built with all the Phase-1 samples and selected edges derived from the edgewise permutation test. At Phase 2, we generalized the prediction model to various independent HCP data and sub-populations, including different task states, gender, and age groups to estimate the in-sample generalization performance. Another two independent datasets were used to assess out-of-sample model generalization (the bottom box). We randomly selected 90% of subjects to calculate generalization performance and this procedure was repeated 10,000 times for the non-parametric test against null distribution. Iter = iteration.

We employed the permutation and bootstrapping procedures to evaluate the statistical significance and reliability of the predictive models. In the bootstrapping procedure, we repeated the 10-fold cross-validation (CV) procedure 1,000 times. Each repetition would result in a slightly different prediction performance *r*_[observed, predicted]_ due to the sampling differences in training and testing sets. Therefore, a prediction performance distribution was generated after 1,000 iterations. To test whether the prediction performance of the model occurred by chance, we adopted the permutation procedure where we scrambled all subjects’ SP scores and their functional connectivity. Each feature and SP scores were permuted independently to generate a fully randomized data matrix. The 10-fold cross-validation procedure with the randomized matrix was repeated 1,000 times to generate null distribution. Statistical significance of the predictive models was determined by comparing the medium of the bootstrapping distribution with the permutation-based null distribution and the 95th percentile points of each distribution were used as the critical values for a one-tailed test against the null hypothesis with *P* = 0.05. With the permutation and bootstrapping procedures, we obtained a set of most frequently selected edges based on the edge-level non-parametric permutation test (*P* < 0.001). We then built prediction models with all the model construction population and the selected edges (positive- and negative-correlation edges separately) for model generalization estimation. We constructed 100 models for generalization where each model was built with a CV of 1,000 iterations (see next session).

### Model generalization and evaluation (Phase 2)

At the model generalization phase, data with different fMRI states (i.e., resting and other task states) and subpopulation groups (i.e., age and gender) were selected from another half of the HCP sample to examine to which extent these factors influence the model generalization. To this end, we randomly selected 150 subject’s resting-state and task-state (including seven tasks) data as generalization sets. The models constructed in phase 1 were then applied to the generalization sets individually to estimate generalization prediction. This in-sample generalization procedure was repeated 1,000 times to obtain a generalization prediction distribution for each model built in phase 1. Similarly, to estimate to which extent the population factors modulate the generalization prediction, we randomly selected 150 females and males as well as 150 adults younger than 30-years-old and adults older than 30-years-old separately. The same generalization procedure was applied to these sub-populations. The Pearson correlation between observed and predicted SP scores was used to assess model generalizability. To minimize sampling bias, the HCP sample splitting procedure (i.e., randomly splitting the HCP into construction and generalization samples) was repeated 100 times. A model was constructed for each split; therefore, 100 models were built at phase 1. As a result, the iteration for cross-validation (phase 1) and generalization (phase 2) procedures was 100,000 times.

To estimate the out-of-sample model generalization performance, we used all the HCP data to build a model and generalized the model to the two independent datasets, i.e., the SLD and ASC. Both datasets are previously published, and they were collected by different research groups with different task settings. We trained this predictive model with all the HCP samples and the FCs of the significant edges in the language task (due to the language task data yielded the best in-sample generalization performance). We then applied this model to the two datasets to generate predicted semantic scores. Pearson correlations between observed and predicted semantic scores were used to assess out-of-sample generalizability. We repeated this out-of-sample generalization and the corresponding permutation procedures 10,000 times (with 90% of the samples each). Statistical significance was determined by the comparison between the bootstrapping distribution and the permutation-based null distribution. All the prediction analyses were performed with customized MATLAB scripts.

### Predictive modeling with different modules intra- and inter-networks connections

To further estimate the prediction contributions of different connection modules, we conducted the predictive modeling with both bootstrapping and permutation procedures for each pair of sub-network connections. First, we classified the semantic network nodes into three sub-networks based on previous resting-state connectivity patterns (Xu *et al*. 2016). The three sub-networks include the Perisylvian network (PSN), frontoparietal network (FPN), and default mode network (DMN). We then separated the connections into six divisions based on their intra- and inter-network connectivity. Specifically, the six divisions (i.e., connection modules) include three modules of intra-network connections (PSN-PSN, FPN-FPN, and DMN-DMN) and three modules of inter-network connections (PSN-FPN, PSN-DMN, and FPN-DMN). The same predictive modeling procedures were conducted for each connection module.

### Control analyses

We further applied our built SP models to predict CC and MC scores to examine whether the SP prediction model is component-specific in explaining individual differences of SP or domain-general that the models can be used to predict other cognitive traits (i.e., CC and MC) that do not require manipulation and processing of semantic information. Moreover, we tested whether a domain-general model can be used to predict SP scores. To do so, we constructed two prediction models for the CC and MC scores with half of the HCP samples and then generalized the two models to predict SP scores in another half of the HCP samples.

## Results

### The core component of semantic processing (SP)

The confirmation factor analysis (CFA) was used to extract a latent variable to reflect SP ability from five behavioral variables (see Fig. 2A for the distributions of the scores of the five tests). Another two latent variables for cognitive control (CC) and motor control (MC) were also estimated for comparisons and control analyses. The three-factor CFA model has a robust model fit to the data (*χ^2^*_(41)_ = 463.264, *P* = 1.00×10^-4^, *RMSEA* = 0.109, *CFI* = 0.750). The SP latent variable explains most of the variances from the five predefined semantic tests (Fig. 2B, left panel). In particular, the picture vocabulary (PicVoc: *R^2^*_PicVoc_ = 0.755) and word reading recognition (ReadEng: *R^2^*_ReadEng_ = 0.841) tests contributed largely to the SP latent scores. The SP latent variable shows a moderate-to-low correlation with the other two latent variables (*R^2^*_[SP,CC]_ = 0.169; *R^2^*_[SP,MC]_ = 0.118; see Fig. 2B, right panel). The distribution of the SP scores across the entire HCP population was shown in Fig. 2C. These SP scores were used for predictive modeling and generalization estimation.

**Fig. 2.**
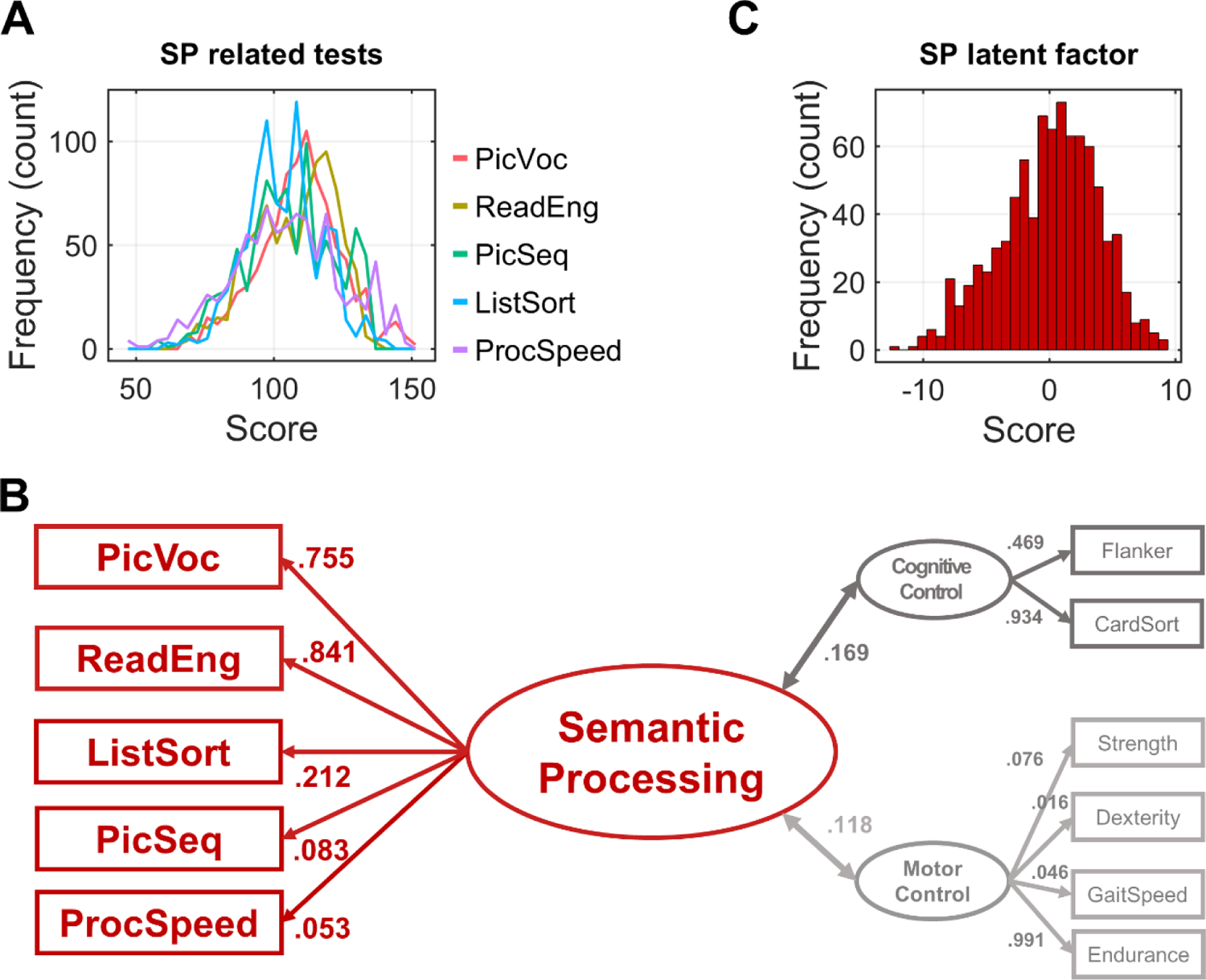
Confirmatory factor analysis (CFA) was used to estimate the latent semantic processing (SP) component and another two general control components. **A**, distributions of five SP-related test variables (see details of the tests and abbreviations in Table S1). **B**, a three-factor CFA model (SP, CC, and MC components) can significantly explain observed test variables. The number beside each line denotes the contribution (*R*^2^) of each test variable to explaining the latent variables. **C**, distribution of the latent SP scores was displayed in the histogram.

### Cross-validated model performance in predicting latent SP scores

At the model construction phase, the SP prediction models built with functional connectivity (FC) features were significantly predictive of individual SP scores (see Fig. 3A left panel for the predicted vs. observed scores). This cross-validation (CV) prediction was not only statistically significant but also reliable across repetitions, which was demonstrated by the bootstrapping and permutation procedures with 100,000 iterations (the positive-predictive model: *r*_pos[obesered, predicted]_ = 0.322, *P* = 2.00×10^-5^; the negative-predictive model: *r*_neg[obesered, predicted]_ = 0.308, *P* = 2.00×10^-5^; Bonferroni-corrected) (Fig. 3A, middle panel). The CV model performance of the language task (LT) data outperformed models of the resting state and other tasks (LT vs. resting state [RS]: *P* = 0.005; LT vs. relational task [RT]: *P* = 0.005; LT vs. gambling task [GT]: *P* = 7.07×10^-5^; LT vs. emotion task [ET]: *P* = 7.03×10^-5^; LT vs. working memory task [WM]: *P* = 0.014; LT vs. motor task [MT]: *P* = 0.153; LT vs. social task [ST]: *P* = 0.191; non-parametric tests with Bonferroni corrected *P*) (Fig. 3A, right panel).

**Fig. 3.**
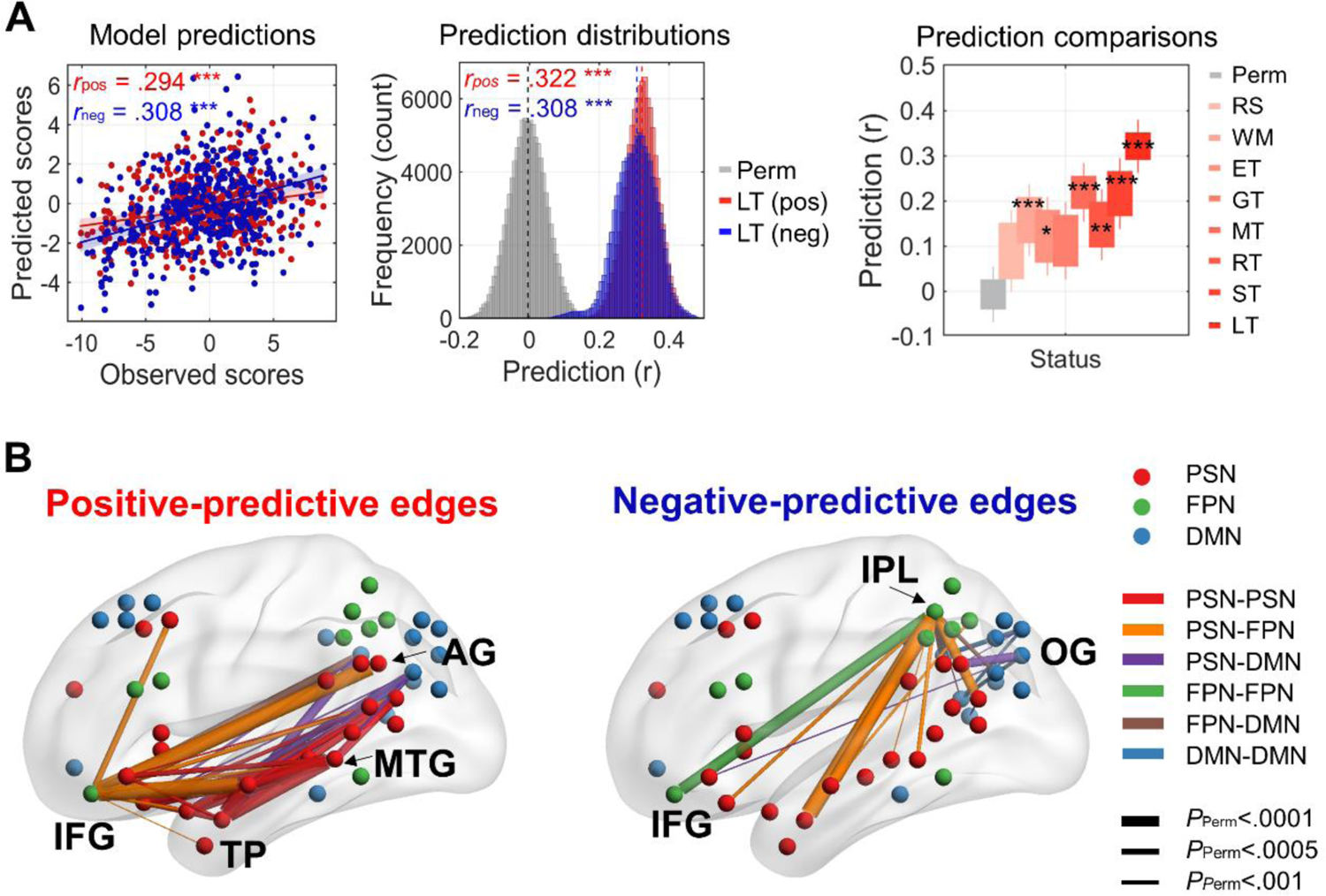
Cross-validated model prediction performance and the significantly contributing connections. **A**, SP prediction performance at the model construction phase. Left panel, scatter plot shows linear correlations between predicted and observed SP scores; Middle panel, bootstrapping- and permutation-based prediction distributions derived from the positively (red) and negatively predictive (blue) FCs, respectively, with language task (LT) data. Perm = permutation; Right panel, prediction distributions across resting and task states. Each box in the boxplot represents the quartile of each prediction distribution. Permutation test: *, *P* < 0.05; **, *P* < 0.01; ***, *P* < 0.005, Bonferroni corrected. Data abbreviations: RS, resting state; WM, working memory task; ET, emotion task; GT, gambling task; MT motor task; RT relational task; ST, social task; LT language task. **B**, significantly predictive edges contributing to the model prediction. The thickness of the connections denotes the statistical significance derived from the edgewise permutation test. Left panel, significant positive-predictive edges; right panel, significant negative-predictive edges. Node abbreviation: IFG, Inferior Frontal Gyrus; TP, Temporal Pole; AG, Angular Gyrus; MTG, Middle Temporal Gyrus; IPL, Inferior Parietal Lobe; OG, Occipital Gyrus.

With the edgewise permutation test, we identified connections (i.e., edges) that were significantly contributing to the CV model predictions. We classified predictive edges into two types (i.e., positively and negatively predictive edges) for the model construction and validation. The positively predictive edges indicate that increased FC strengths are associated with enhanced SP ability. These positively predictive edges are mainly those PSN intra-network edges and PSN-FPN and PSN-DMN inter-network edges (see Fig. 3B, left panel), particularly including the edges between the orbital part of the inferior frontal gyrus (IFG), temporal pole (TP), middle temporal gyrus (MTG), and angular gyrus (AG). In contrast, the negatively predictive edges indicate that increased FC strengths are associated with poorer SP ability. We found that the negatively predictive edges distributed across connection modules, mainly between an FPN node, inferior parietal lobule (IPL), and PSN temporal nodes, as well as intra-FPN connections (e.g., an edge between IFG and IPL) (Fig. 3B, right panel). All the significantly contributing edges were listed in Table S3.

With predictive modeling for each module of connections (i.e., PSN-FPN, PSN-DMN, PSN-PSN, FPN-DMN, DMN-DMN, and FPN-FPN; see Fig. 4A for the edge patterns), we found that different modules showed different degrees of predictive power (Fig. 4B). The significant positively predictive edges are those module connections between PSN and the other two sub-networks as well as the PSN intra-network connections (*r* _PSN-FPN_ = 0.220, *P =* 6.00×10^-5^; *r* _PSN-DMN_ = 0.220, *P =* 6.00×10^-5^; *r* _PSN-PSN_ = 0.219, *P =* 1.52×10^-4^; *r* _FPN-DMN_ = 0.176, *P* = 0.002; *r* _DMN-DMN_ = 0.154, *P* = 0.007; *r* _FPN-FPN_ = 0.075, *P =* 0.300; Bonferroni corrected *P*; Fig. 4B). The significant negatively predictive edges are mainly those between PSN and DMN as well as connections between PSN and FPN (*r* _PSN-FPN_ = 0.235, *P =* 6.00×10^-5^; *r* _PSN-DMN_ = 0.199, *P =* 4.80×10^-4^; *r* _PSN-PSN_ = 0.066, *P =* 0.587; *r* _FPN-DMN_ = 0.223, *P* = 6.09×10^-5^; *r* _DMN-DMN_ = 0.203, *P* = 3.06×10^-4^; *r* _FPN-FPN_ = 0.151, *P =* 0.007; Bonferroni corrected *P*) (Fig. 4B). Detailed predictive edges for each module were displayed in Fig. 4C. These module-based prediction results are consistent with the overall edgewise prediction results described in Fig. 3B.

**Fig. 4.**
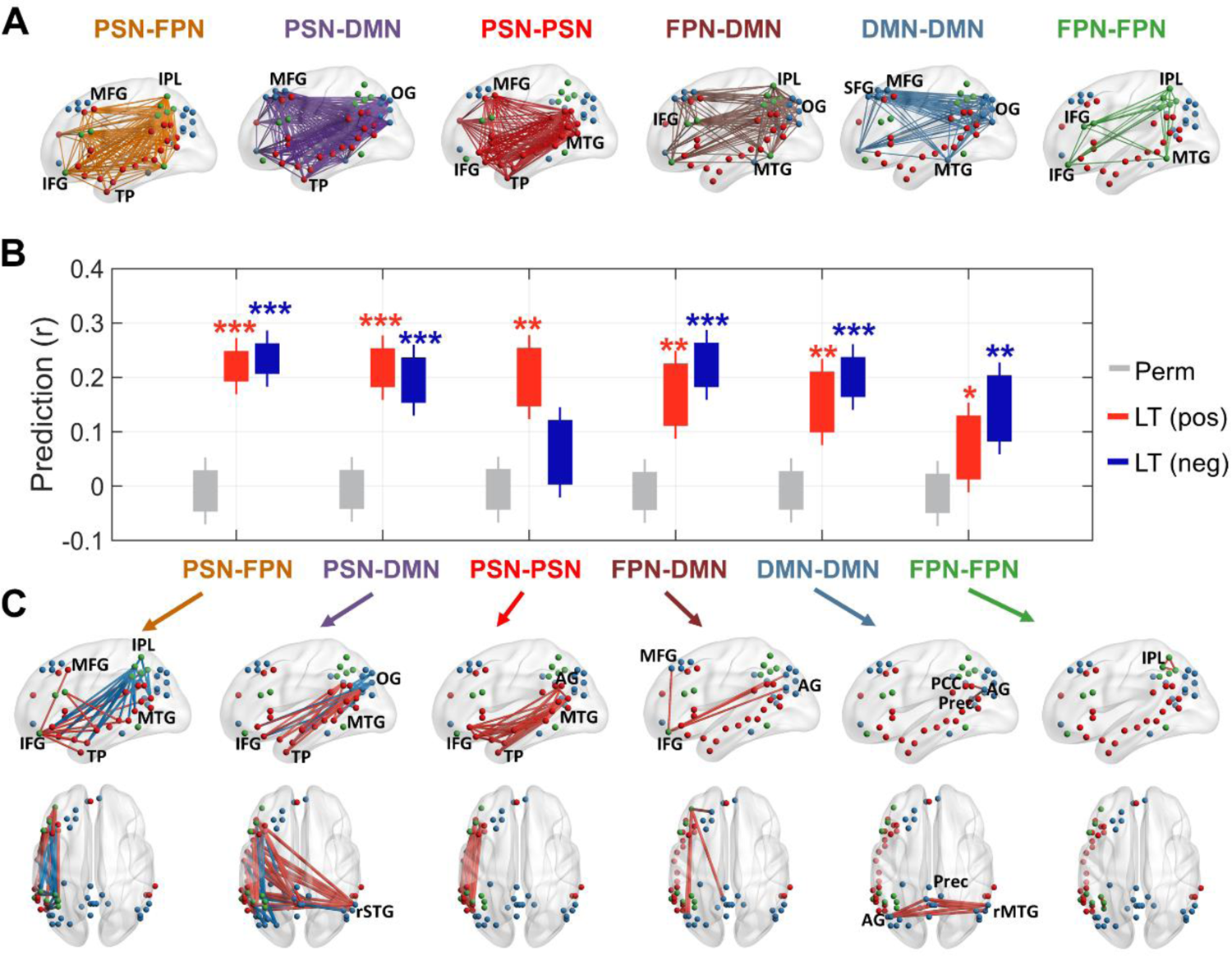
Predictive modeling (Phase 1) for each module of connections. **A**, six connection modules. Key regions in each module were labeled. **B**, model prediction powers for each module and each type of edge. The red and blue boxes represent the predictive power of positive- and negative-predictive models, respectively. The order of the modules was sorted by the mean prediction *r* value of the positive models in descending order. Perm, permutation. **, *P* < 0.01; ***, *P* < 0.005, Bonferroni corrected. **C**, significantly predictive edges for each connection module (edgewise permutation test, *P* < 0.005). The red- and blue-color edges represent the significant positively and negatively predictive connections, respectively. Node abbreviation: IFG, Inferior Frontal Gyrus; TP, Temporal Pole; AG, Angular Gyrus; MTG, Middle Temporal Gyrus; IPL, Inferior Parietal Lobe; OG, OccipitalGyrus; MFG, Middle Frontal Gyrus; PCC, Posterior Cingulum Cortex; Prec, Precuneus; rSTG, right Superior Temporal Gyrus; rMTG, right Middle Temporal Gyrus.

### Model generalization in predicting independent samples

To estimate model generalization, we constructed predictive models with all the data used at Phase 1 and the significant edges derived from the edgewise permutation test (*P* < 0.001). We used the language task’s fMRI data to build the models for generalization because the CV performance of the language task data outperforms the resting state and other tasks (see Fig. 3A, the right panel). The in-sample and out-of-sample generalization performances of the predictive models were assessed with another half of the unseen HCP dataset and the two independent datasets, respectively. For in-sample generalization, different task states (i.e., resting-state and seven tasks) and sub-populations (i.e., two age groups and two gender groups) of the HCP dataset were used to estimate to which extent these variables affect the generalization performance. First, we generalized our trained model to the data derived from different states. We found that positively predictive model significantly generalized to unseen individuals for the language task data (LT: median *r* = 0.249, *P* = 0.0068; permutation test; the same for the following tests) and the motor task (MT: *r* = 0.208, *P* = 0.037), but not for the other states (RS: *r* = 0.105, *P* = 0.792; WM: *r* = 0.157, *P* = 0.200; ET: *r* = 0.132, *P* = 0.405; GT: *r* = 0.139, *P* = 0.341; RT: *r* = 0.092, *P* = 0.999; ST: *r* = 0.154, *P* = 0.226; Bonferroni corrected *P*). The generalization performance of the language task was significantly better than the resting state (*P* = 0.016; uncorrected). However, no significant difference was found between the language task and most of the other task states in generalization (LT vs. WM: *P* = 0.090; LT vs. ET: *P* = 0.044; LT vs. GT: *P* = 0.055; LT vs. MT: *P* = 0.188; LT vs. RT: *P* = 0.017; LT vs. ST: *P* = 0.067; uncorrected). These results indicated that the language task selectively enhances the model generalization (Fig. 5A, left panel), which consistent with the cross-validation results described above. To further explore which module of connections contributing to the task effect in generalization, we conducted the model generalization analysis for different intra- and inter-network connections. We found that only those edges connecting to the PSN yielded significant model generalization with language task data (*r* _PSN-PSN_ = 0.248, *P =* 0.005; *r* _PSN-FPN_ = 0.239, *P =* 0.007; *r* _FPN-DMN_ = 0.199, *P =*0.020; Bonferroni corrected *P*). While the PSN intra-network connections (i.e., PSN-PSN) contribute most to the task enhancement in generalization, their model generalization in the language task also outperformed that in the resting state (LT vs. RS: *P =* 0.039, uncorrected; Fig. 5A, right panel).

**Fig. 5.**
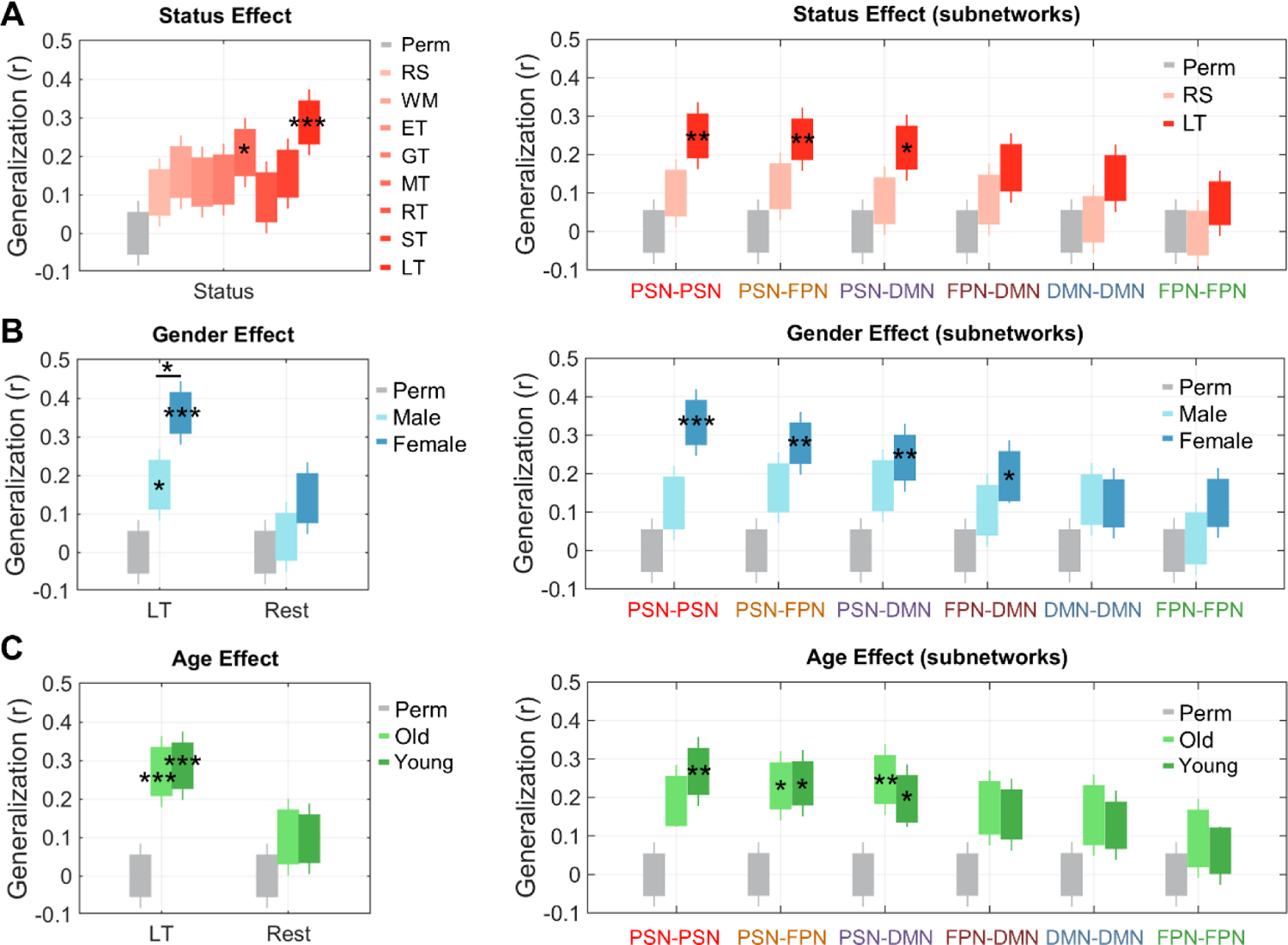
The in-sample generalization performance of the positive predictive models was modulated by task and gender. **A**, model generalization performance with independent data across task states. Abbreviation: perm, permutation; RS, resting state; WM, working memory task; ET, emotion task; GT, gambling task; MT, motor task; RT, relational task; ST, social task; LT, language task. **B**, predictive model generalization performance with independent data of the two gender groups. LT, language task; Rest, resting state. **C**, model generalization performance for the two age groups. Right panels in A-C show generalization evaluation for the six connection modules for the status, gender, and age effect, respectively. These connection modules are sorted according to the significance of the status effect in descending order. Boxplots represent 25th and 75th percentiles (box) and range (whiskers) for the distributions. Asterisks indicate the significance of the generalizability: *, *P* < 0.05; **, *P* < 0.01; ***, *P* < 0.005; Bonferroni corrected.

We further examined the in-sample generalization performance to different gender and age groups of the HCP dataset with the positive predictive models (see Fig. 5B&C). We selected the resting state and language task data of these sub-populations as the target data for generalization. We examined to which extent these population variables modulate the generalization performance. For the language task data, the trained model significantly generalized to both females (*r* = 0.359, *P* = 2.00×10^-5^, Bonferroni corrected) and males (*r* = 0.174, *P* = 0.030; Bonferroni corrected). The model’s generalizability was significantly higher for females than males with the language task data (*P* = 0.039; Bonferroni corrected) (Fig. 5B, left panel). Those edges connecting to the PSN nodes yielded significant model generalization only for females (*r* _PSN-PSN_ = 0.332, *P =* 1.20×10^-4^; *r* _PSN-FPN_ = 0.279, *P =* 0.002; *r* _PSN-DMN_ = 0.240, *P =* 0.009; *r* _FPN-DMN_ = 0.190, *P =* 0.045; Bonferroni corrected). The gender effect in generalization was most prominent in the PSN intra-network connections (males vs. female: *P* = 0.014, uncorrected) (Fig. 5B, right panel), which is similar to the task effect. No significant generalization prediction was observed for resting-state data for both gender group (female: *r* = 0.140, *P* = 0.080; male: *r* = 0.040, *P* = 0.624; corrected). We further tested whether the homogeneity in FC strengths within the females was significantly less than that within the males. We randomly selected 100 males and females respectively and calculated their standard deviations (*SD*; a proxy of inhomogeneity) of the FC strengths of the significantly predictive edges (displayed in Fig. 3B). This process was repeated 10,000 times to estimate *SD* distributions. We found that the *SD* of the FC for the females was significantly smaller than that for the males (*P* = 0.001, non-parametric test), which suggests that females’ SP-related FCs are more homogeneous than males. Moreover, for both age groups, significant generalization predictions were found only for the language task data (young: *r =* 0*.2*85, *P =* 2.60×10^-4^; old: *r =* 0.269, *P =* 5.80×10^-4^, corrected), consistent with the overall generalization results. However, no significant difference in generalization was found between the younger and older participants, neither for the language task (*P =* 0.874; corrected) nor for resting-state data (*P =* 0.873; corrected) (Fig. 5C).

For the generalizability of the negative-predictive models, similar task status and gender effects were found as the positive-predictive models (Fig. S1). First, significant generalization was found only for the LT data (*r* = 0.316, *P* = 3.20×10^-4^, Bonferroni corrected), but not for any of the other states (*P*s > 0.1). The generalization performance of the language task data was significantly better than the resting state (*P* = 0.014, Bonferroni corrected) and most of the other tasks (LT vs. ST: *P* = 0.018; LT vs. ET: *P* = 0.037; LT vs. GT: *P* = 0.070; LT vs. MT: *P* = 0.041; LT vs. RT: *P* = 0.039; LT vs. WM: *P* = 0.160; Bonferroni corrected; see Fig. S1A, left panel). We further found that most of the subnetwork connections (i.e., PSN-FPN, PSN-DMN, DMN-DMN, FPN-DMN, and FPN-FPN) except the PSN intra-network connections (i.e., PSN-PSN) contributed to the task enhancement in model generalization (LT vs. RS: *P* _PSN-FPN_ *=* 0.023, *P* _PSN-DMN_ *=* 0.013, *P* _DMN-DMN_*=* 0.024, *P* _FPN-DMN_ *=* 0.016, and *P* _FPN-FPN_ *=* 0.031; uncorrected) (Fig. S1B, right panel). However, these comparisons were not survived after correction (LT vs. RS: *P* _PSN-FPN_ *=* 0.138, *P* _PSN-DMN_ *=* 0.075, *P* _DMN-DMN_ *=* 0.142, *P* _FPN-DMN_ *=* 0.096, and *P* _FPN-FPN_ *=* 0.183). For the generalization in different gender and age groups, the model’s generalizability was significantly higher for females than males only for the language task data (*P* = 0.043, uncorrected) (Fig. S1B, left panel). However, this gender effect in generalization did not reach significance in any module of the connections (Fig. S1B, right panel), but the quantity differences between females and males were observed across modules except PSN’s intra-network connections. Also, no significant difference in generalization was found between the young and old participants, neither for the language task (*P =* 0.660, corrected) nor resting-state (*P =* 0.495, corrected) data (Fig. S1C).

### Out-of-sample model generalization evaluation

To further examine to which extent the cross-validated models can be generalized to independent datasets (i.e., out-of-sample generalization) collected in different populations and with different task settings and SP measures, we used our HCP model to predict another two dataset’s semantic measures: a classic semantic lexical decision task and a natural story comprehension task. The two tasks were chosen because they represent classic semantic processing paradigms with and without experimental control. The out-of-sample model generalization was often unsuccessful in previous attempts. Therefore, we consider this analysis as an exploratory test and selected a wide range of percentages of significant edges as features to build HCP models to examine whether the number of connections modulates the generalization. We found that only the positive-predictive model can significantly predict unseen individuals’ reading comprehension scores in the ASC dataset across a wide range of edge selection percentages (e.g., 50% edge: *r* = 0.491, *P* = 0.005, uncorrected) (Fig. 6A). However, the positive HCP model can only predict semantic lexical decision (SLD) scores when 50% of the predictive edges were used (*r* = 0.350, *P* = 0.047, uncorrected). No significant out-of-sample generalization was found when we applied the negative-predictive models (Fig. 6B). We also tried to combine both the positive and negative models for generalization. However, the model generalization performance did not outperform the positive-predictive model alone.

**Fig. 6.**
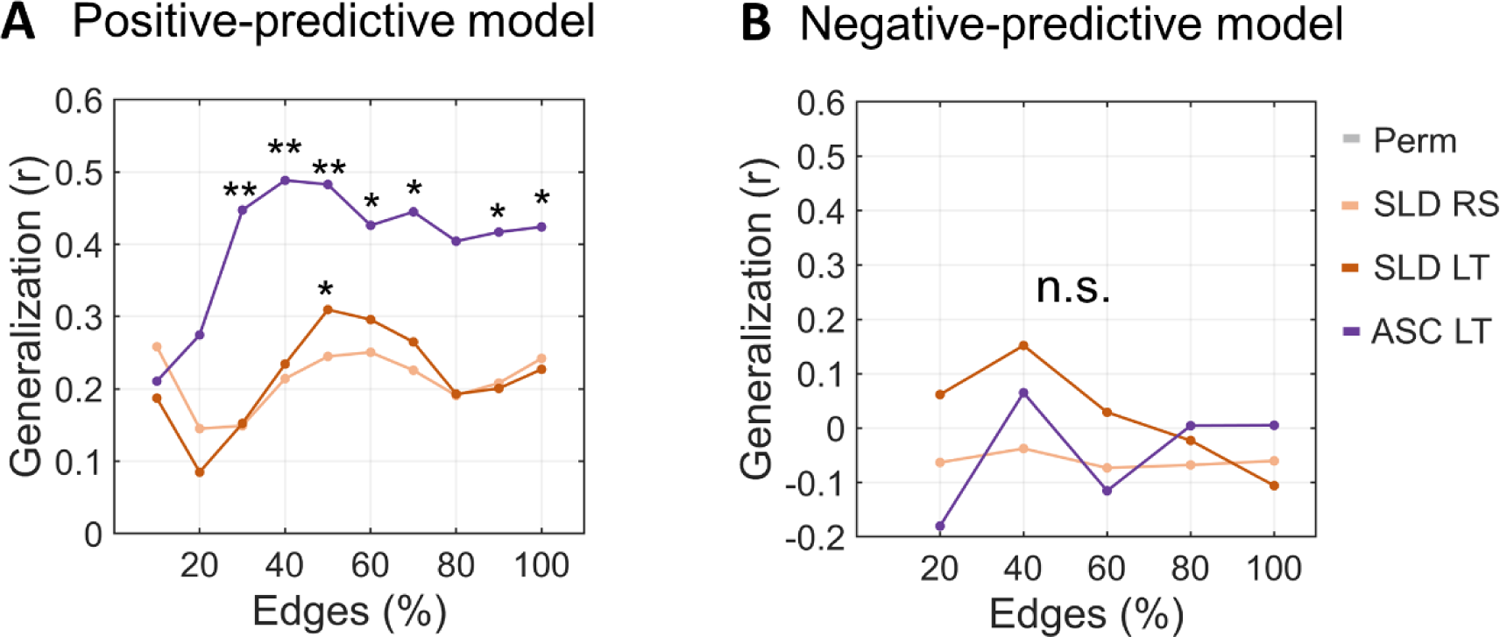
Out-of-sample generalization performance with two independent datasets and semantic measures. **A**, the out-of-sample generalization of the positive-predictive model. The line chart illustrates the out-of-sample generalization performance using different percent of predictive edges. **B**, the out-of-sample generalization performance of the negative-predictive model. Dataset abbreviation: SLD RS, resting-state data from the SLD dataset; SLD LT, task-state data from the SLD dataset; ASC LT, language task data from the ASC dataset. Edges (%), percentage of significant edges that used to build an HCP model for generalization. *, *P* < 0.05; **, *P* < 0.01; uncorrected.

### Control analyses

To further examine to which extent the HCP SP model is component-specific (selectively predicting SP scores), we used the built SP model to predict CC and MC scores. We found that the positive predictive model was not predictive of CC nor MC while the negative-predictive model was weakly predictive of CC but was not predictive of MC (Fig. S2A, *SI*). The results indicate that the SP model is largely specific to predict SP ability. In addition, we further examine whether the CC and MC models built with half of the HCP data can be used to predict SP scores of another half of the data. We ran the predictive modeling analysis (including the phase 1 model construction and phase 2 generalization) with the CC and MC scores. The models built with the CC and MC scores cannot significantly predict held-out subjects’ SP scores (Fig. S2B&C, *SI*). These findings provide converging evidence in support of the specificity of the SP model and the FC patterns underlying individual differences in SP.

## Discussion

The present study used connectome-based predictive modeling and datasets with sizable samples to examine the functional network organizations underlying individual differences in semantic processing (SP) ability. We constructed predictive models with rigorous cross-validation, bootstrapping, and permutation procedures and assessed the model’s generalizability with unseen in-sample and out-of-sample datasets. We demonstrate the robust relationships between individual differences in SP and variabilities in functional connectivity (FC) patterns while overcoming the low effect size inherent by small sample sizes and traditional correlational approaches. We also identified a cluster of intra- and inter-network FCs where their variabilities across subjects contributed significantly to predict the latent SP scores. Increased FCs within the Perisylvian network (PSN) and between PSN and other sub-networks are predictive of superior SP ability, while increased FCs between a frontoparietal network (FPN) node, inferior parietal lobe, and other sub-networks are predictive of poorer SP. These predictive relationships were enhanced when subjects participated in a language task comparing to resting-state and other tasks. This task enhancement in prediction is more prominent for females than males. Also, the SP prediction model built with all HCP data can be generalized to independent datasets that used very different neuroimaging and behavioral paradigms. These findings — that complex FC models predict different measures of SP across different populations — not only provide significant insights into our understanding of the neural network organizations underlying individual differences in SP but also further provide detailed effects in how task and demographic factors modulate the neural-SP relationships.

The current models, which are constructed by the HCP dataset and capable of generalizing to different in-sample and out-of-sample datasets and SP behavioral measures, make significant progress toward identifying neuromarkers of SP. SP is not a unitary ability, rather, it has been proposed to be related to various cognitive and control components. Therefore, it is difficult to select an unbiased behavioral task that can reflect the core composition of SP while minimizing the task-specific components so that the built model can be generalized to other semantic task contexts. Here we selected five different behavioral measures and used confirmation factor analysis to extract a latent SP component, which could minimize biases in task selection and ensure representation of the SP measure. Also, we included two domain-general latent factors, cognitive control (CC) and motor control (MC) where their test materials do not involve any semantic stimuli and their tasks are not required any SP processes. Additional control analyses confirmed that neither the CC nor the MC model was predictive of SP scores, while the SP models were not consistently predictive of CC or MC scores neither, which suggest that the current SP models and the underlying connectivity organization are largely specific to the SP component that require the manipulation and process of semantic or conceptual information.

In addition to demonstrating the prediction performance of the SP model, we further reveal that FCs connecting to the PSN nodes play a critical role in the model prediction and generalization, especially the interplay between the FPN and PSN. Increased FCs within PSN and between PSN and FPN is associated with superior SP ability, especially between the inferior frontal gyrus (IFG) and distributed temporoparietal PSN nodes. Previous studies have shown that both structural and functional connectivity properties of the PSN were associated with semantic task performance (Bookheimer, 2002; Saur *et al*., 2008). The predictive PSN intra-network connections mainly included the connections between the left middle temporal regions (e.g., anterior to posterior) and the left IFG. These regions are the main constitutes of the canonical language system that is more activated by language tasks compared with control tasks (Fedorenko *et al*., 2011; Fedorenko & Thompson-Schill, 2014). For example, the left middle temporal gyrus (LMTG) has been proposed to acting as a hub in the semantic and language network, where it has widely distributed connections with other language areas (Turken & Dronkers, 2011). The FCs between the LMTG and other regions, including the left IFG, dorsal lateral and medial prefrontal cortex are associated with individual differences in semantic behaviors (Jackson *et al*., 2016; Wei *et al*., 2012). Our results are also consistent with previous studies on aphasia patients and stroke patients, where a selective disruption of IFG and connections between IFG and left anterior superior temporal regions was associated with semantic impairments (Meinzer *et al*., 2011). The predictive FCs in PSN and FPN identified by the current study could potentially be neural indicators of language/semantic impairments/disorders and neural predictors for future intervention since these connections have been demonstrated to be tightly linked to language functions (e.g., semantic and syntactic processes) (Badre *et al*., 2005; Krieger-Redwood *et al*., 2016; Papoutsi *et al*., 2011; Snijders *et al*., 2010; Tyler & Marslen-Wilson, 2007; Vatansever *et al*., 2017).

We also found that increased several network FCs within FPN and between FPN, default mode network (DMN), and PSN are predictive of poorer SP ability. One critical node in FPN, the left inferior parietal lobule (LIPL) is significantly contributed to the negative predictions. Increased FC strengths between the LIPL and a range of PSN nodes across the inferior frontal and middle temporal regions are associated with decreased SP ability. LIPL has been characterized as a provincial hub and inter-network connector connecting FPN with both PSN and DMN. This region has been proposed to be connecting the semantic control system in FPN with a putative language-based semantic system in PSN and a memory-based simulation system in the DMN (Xu *et al*., 2016). Increased connectivity of the LIPL with the other two systems may be related to decreased efficiency in semantic processing in a semantic task. For example, in a challenging semantic retrieval task, people with superior SP ability can solve the task relatively fast and efficiently where they may only rely on increased focal FCs between the frontotemporal PSN regions. However, people with poor SP ability may need additional assistance from other semantic systems, especially the FPN control system to solve the difficult task and the memory-based system in DMN for retrieving additional information. One interpretation of the involvement of LIPL connections is that increased FCs between PSN regions and the inter-network connector LIPL is a compensation mechanism for those people with poor SP while they need to recruit additional SP-related information to make a decision. This compensation interpretation may also be used to explain why increased FCs between PSN posterior temporal regions and DMN occipital regions are associated with poorer SP ability. Future studies should explore and test this possibility.

We further demonstrate that the built SP models can be generalized to independent unseen in-sample and out-of-sample data. We explore to what extent the task and demographic factors (i.e., gender and age) modulate the generalization performance. We found that both task and gender are two critical factors affecting how well a trained model generalizing to unseen subjects. Language task significantly enhances the generalization performance over resting-state and other task-state data, which is consistent with the prediction performance in the cross-validation. This language task enhancement effect is consistent with previous findings from group-level FC studies where they demonstrated increased FCs between the frontoparietal cognitive-control regions and language areas when subjects performed language tasks (Cole *et al*., 2014; Di *et al*., 2013; T. Jiang *et al*., 2004; Smith *et al*., 2009). For individual prediction findings, previous studies show that cognitive tasks enhance individual differences of fluid intelligence in FC patterns, such that predictive models built from task fMRI data outperform models built from resting-state fMRI data (Greene *et al*., 2018; R. Jiang *et al*., 2020). Consistent with and moving beyond this finding, here we demonstrate that only language tasks can amplify individual differences in FC patterns that are more tightly related to individuals’ behavioral semantic performance than other tasks. We demonstrate that this language task amplification effect can be observed not only for cross-validation prediction but also for model generalization. One possibility of this task-amplification in model prediction is that the language task-induced functionally relevant FC changes over resting state and other task states subserving performance of the semantic tasks not only at the group level but also at individual differences level. During a language task, the FCs reorganize based on the task demand where the FC organization is optimal for processing the language stimuli in hand (Feng *et al*., 2015), especially semantic information. Therefore, the individual differences in FC during language tasks are best associated with behavioral test scores that involve similar semantic processes, whereas resting-state FCs are unconstraint, and it is likely to involve many SP-irrelevant components, e.g., mind wandering (Godwin *et al*., 2017), arousal (Koike *et al*., 2011), attention (Bonnelle *et al*., 2011), and different levels of conscious thoughts (Smallwood *et al*., 2012).

In addition to the task effect, we also demonstrate gender differences in model generalization. We found that model generalization to females was more robust than males. This may be due to gender differences in SP-related FC organization, task-induced neural activation patterns, or both (Baxter *et al*., 2003; Biswal *et al*., 2010; Satterthwaite *et al*., 2015; Scheinost *et al*., 2015). For example, previous studies have revealed females had more focal activation in the left hemisphere and greater right posterior temporal and insula SP-related activations than males (Baxter *et al*., 2003). In FC patterns, detectable gender differences were found across network modules and seed-based connectivity strengths (Biswal *et al*., 2010). Consistent with these findings, recent FC studies demonstrate that multivariate resting-state FC patterns are associate with individuals’ cognitive profiles of “male” and “female” (Satterthwaite *et al*., 2015). Extending these previous findings, we demonstrate that model generalization is more robust for females than males. Those sex differences in neurocognitive measures reported in previous studies may not fully explain the underlying cause of the generalization differences found here. One source of the gender differences in model generalization is the differences in FC homogeneity between the two groups. We show that FCs in females are significantly more homogeneous (less inter-subject variability) than that of males. More similar in FC patterns between females than between males at the population level would results in better generalization to unseen females than males. This interpretation and observation suggest that researchers should consider sample homogeneity when building prediction models. The group-specific model may yield superior prediction performance than group-general models. Future studies would need to be conducted to systematically explore this possibility.

Moving beyond cross-validation prediction, we adopted not only in-sample but also out-of-sample generalization estimation with independent datasets. Overestimation of prediction performance is commonly found with traditional correlational approaches and small sample size on one hand while a failure to maintain the independence of training and test datasets in another hand. To ensure data independence, we ensure that model validation and estimation were true tests of the models’ ability to generalize to unseen subjects at every step of the analysis. We also define the predictions based on the levels of generalization to unseen subjects (i.e., level 1: cross-validation; level 2: in-sample generalization; level 3: out-of-sample generalization) with an increasing level of data independence. For prediction with the out-of-sample generalization procedure, the model built with all HCP data can be generalized to unseen datasets with totally different populations, task settings, instruction, data acquisition, MRI scanners, etc. This finding implicates that the SP prediction model captures the core and maybe universal relationships between the variability in SP component and individual variability in FC organization. Nevertheless, there is a limitation of the out-of-sample generalization estimation in the current study. We only included two previously published datasets as the out-of-sample datasets and the sample sizes of the datasets are relatively small, which could potentially limit our examination of the out-of-sample model generalization. Further studies need to be conducted to assess the out-of-sample generalization performance with more and sizable datasets and further examine what factors may modulate the generalization across datasets.

In summary, we have shown that functional connectivity patterns play a critical role in explaining the individual differences in semantic processing ability. The SP prediction model constructed from the HCP dataset can be generalized to independent cohorts with different experimental settings, suggesting high model reliability and generalization. FCs connecting to the Perisylvian language network show the most reliable contributions to predictive modeling and the out-of-sample generalization. These findings contribute to our understanding of the neural sources of individual differences in SP, which potentially lay the foundation for personalized education and improve intervention practice for patients with SP and language deficits.

## Funding

This work was supported by grants from the General Research Fund (Ref. No. 14619518 to Gangyi Feng) by the Research Grants Council of Hong Kong and Direct Grant for Research (Ref. No. 4051137 to Gangyi Feng) by The Chinese University of Hong Kong.

## Acknowledgments

Data collection and sharing for this project was provided by the MGH-USC Human Connectome Project (HCP; Principal Investigators: Bruce Rosen, M.D., Ph.D., Arthur W. Toga, Ph.D., Van J. Weeden, MD). HCP funding was provided by the National Institute of Dental and Craniofacial Research (NIDCR), the National Institute of Mental Health (NIMH), and the National Institute of Neurological Disorders and Stroke (NINDS). HCP data are disseminated by the Laboratory of NeuroImaging at the University of Southern California.

## Potential Competing Interest

Patrick C. M. Wong is a founder of a company in Hong Kong supported by a Hong Kong SAR government startup scheme for universities.

## Code availability

MATLAB scripts to run the main predictive modeling analyses can be found at (https://osf.io/b9h2x/). MATLAB scripts written to perform additional control analyses are available from the authors upon request.

## Data availability

The HCP data that support the findings of this study are publicly available on the ConnectomeDB database (https://db.humanconnectome.org). The meta-data and functional connectivity data can be found at (https://osf.io/b9h2x/).

## Supplementary Information

### Supplementary Methods

#### FMRI procedures in HCP dataset

Resting-state fMRI (1200 frames/scan).

Subjects were asked to lie with eyes open, with “relaxed” fixation on a white cross (on a dark background), think of nothing in particular, and not fall asleep.

#### Working memory fMRI (405 frames/scan)

Participants were presented with blocks of trials that consisted of pictures of places, tools, faces, and body parts (non-mutilated parts of bodies with no “nudity”). Within each run, the four different stimulus types were presented in separate blocks. Also, within each run, ½ of the blocks use a 2-back working memory task, and ½ use a 0-back working memory task (as a working memory comparison).

#### Gambling task fMRI (253 frames/scan)

Participants play a card-guessing game where they are asked to guess the number on a mystery card (represented by a “?”) to win or lose money. Participants are told that potential card numbers range from 1-9 and to indicate if they think the mystery card number is more or less than 5 by pressing one of two buttons on the response box. Feedback is the number on the card (generated by the program as a function of whether the trial was a reward, loss, or neutral trial) and either: 1) a green up arrow with “$1” for reward trials, 2) a red down arrow next to −$0.50 for loss trials; or 3) the number 5 and a grey double-headed arrow for neutral trials.

#### Motor task fMRI (284 frames/scan)

The participants are presented with visual cues that ask them to tap their left or right fingers, squeeze their left or right toes, or move their tongue to map motor areas.

#### Language task fMRI (316 frames/scan)

The language task fMRI scan consists of two runs that each interleaves 4 blocks of a story task and 4 blocks of a math task. The story blocks present subjects with brief auditory stories (5–9 sentences) adapted from Aesop’s fables, followed by a 2-alternative forced-choice question that asks the subjects about the topic of the story. The math task also presents trials auditorily and requires the subjects to complete addition and subtraction problems. The subjects push a button to select either the first or the second answer. For more details about the task, see Binder and Desai (2011).

#### Social-cognition task fMRI (274 frames/scan)

The participants are presented with short video clips (20 s) of objects (squares, circles, triangles) either interacting in some way or moving randomly. After each video clip, the participants chose between 3 possibilities: whether the objects had a social interaction (an interaction that appears as if the shapes are taking into account each other’s feelings and thoughts), Not Sure, or No interaction (i.e., there is no obvious interaction between the shapes and the movement appears in random). Each of the two task runs has 5 video blocks (2 Mental and 3 Random in one run, 3 Mental and 2 Random in the other run) and 5 fixation blocks (15 s each).

#### Relational task fMRI (232 frames/scan)

The stimuli are 6 different shapes filled with 1 of 6 different textures. In the relational processing condition, the participants are presented with 2 pairs of objects, with one pair at the top of the screen and the other pair at the bottom of the screen. They are told that they should first decide what dimension differs across the top pair of objects (shape or texture) and then they should decide whether the bottom pair of objects also differ along that same dimension (e.g., if the top pair differs in shape, does the bottom pair also differ in shape). In the control matching condition, the participants are shown two objects at the top of the screen and one object at the bottom of the screen, and a word in the middle of the screen (either “shape” or “texture”). They are told to decide whether the bottom object matches either of the top two objects on that dimension (e.g., if the word is “shape”, is the bottom object the same shape as either of the top two objects).

#### Emotion task fMRI (176 frames/scan)

The participants are presented with blocks of trials that ask them to decide either which of two faces presented on the bottom of the screen match the face at the top of the screen, or which of two shapes presented at the bottom of the screen match the shape at the top of the screen. The faces have either angry or fearful expressions. Trials are presented in blocks of 6 trials of the same task (face or shape), with the stimulus presented for 2s and a 1 s ITI. Each block is preceded by a 3s task cue (“shape” or “face”) so that each block is 21 s including the cue. Each of the two runs includes 3 face blocks and 3 shape blocks.

**Fig. S1.**
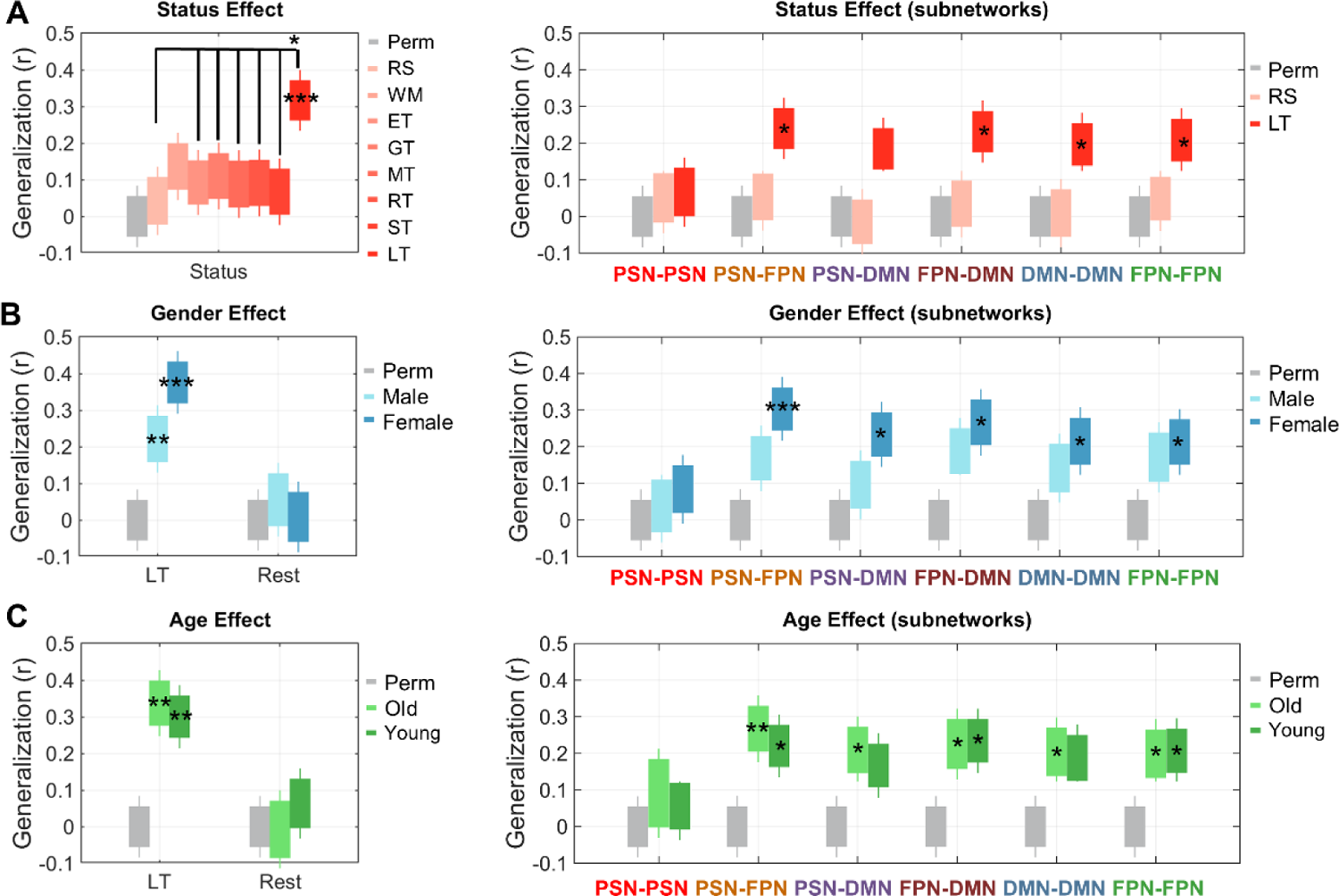
The in-sample generalization performance of the negative-predictive models was modulated differently by task and demographic factors. **A**, model generalization performance with unseen HCP datasets across task states. **B**, model generalization performance with unseen subpopulation for the two gender groups separately. **C**, model generalization performance for the two age groups (> 30-year-old vs. < 30-year-old) separately. Right plots of each panel show the generalization evaluation for each module of connections for the task, gender, and age effect, respectively. Boxplots represent 25th and 75th percentiles (box) and range (whiskers) for all graphs. Asterisks indicate the significance of the generalizability: *, *P* < 0.05; **, *P* < 0.01; ***, *P* < 0.005; Bonferroni corrected.

**Fig. S2.**
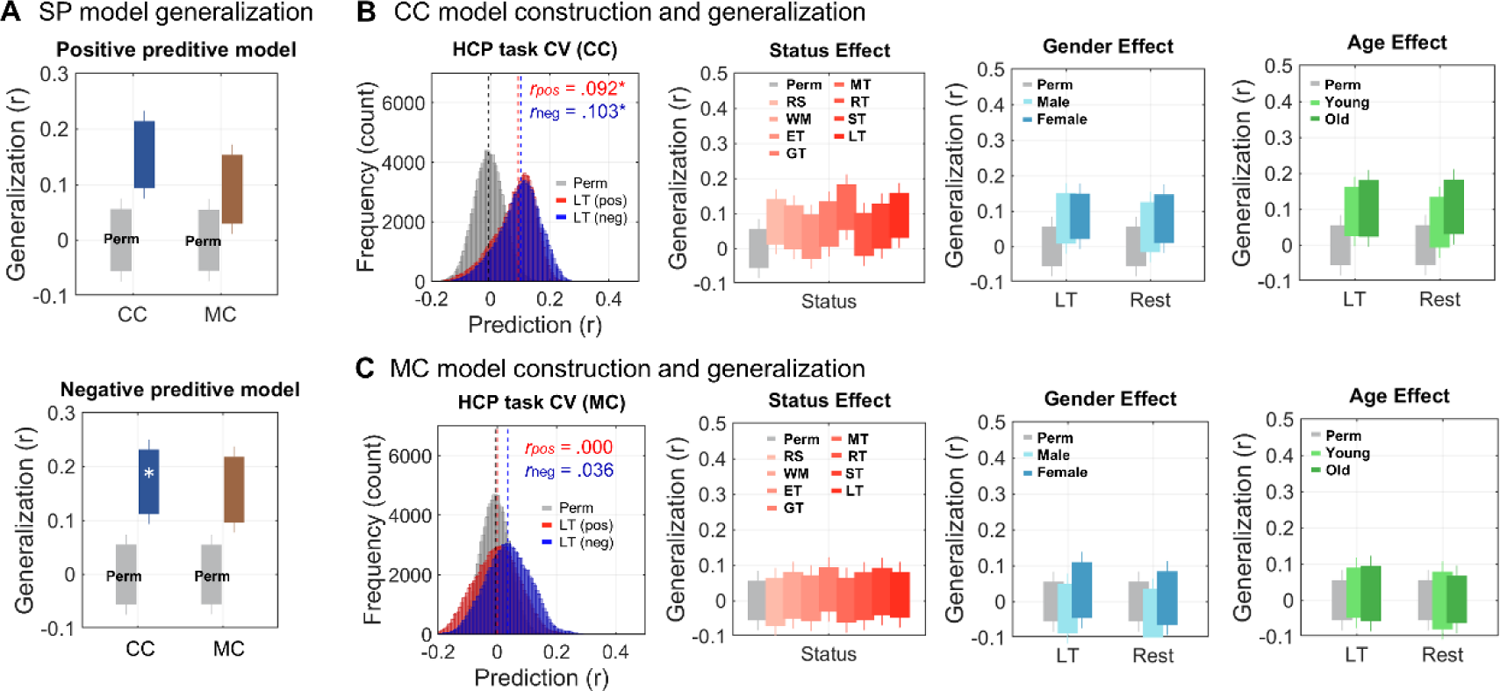
Control analyses. SP model generalization and predictive modeling for the cognitive control (CC) and motor control (MC) components. **A**, SP model’s generalization performance to predict CC and MC scores. *, Bonferroni corrected *P* < 0.05. **B**, prediction performance of the CC model. HCP task CV (CC), cross-validation prediction performance with language task (LT) data. LT (pos) = positive-predictive model; LT (neg) = negative-predictive model; perm = permutation distribution; Status/Gender/Age effect, no significant model generalization to various task statuses, gender and age groups for CC prediction. Abbreviation: Rest/RS = Resting state; WM = working memory task; ET = emotion task; GT = gambling task; MT = motor task; RT = relational task; ST = social task; LT = language task. **C**, model prediction and generalization performance of the MC model. No significant model prediction was found.

**Table S1.**
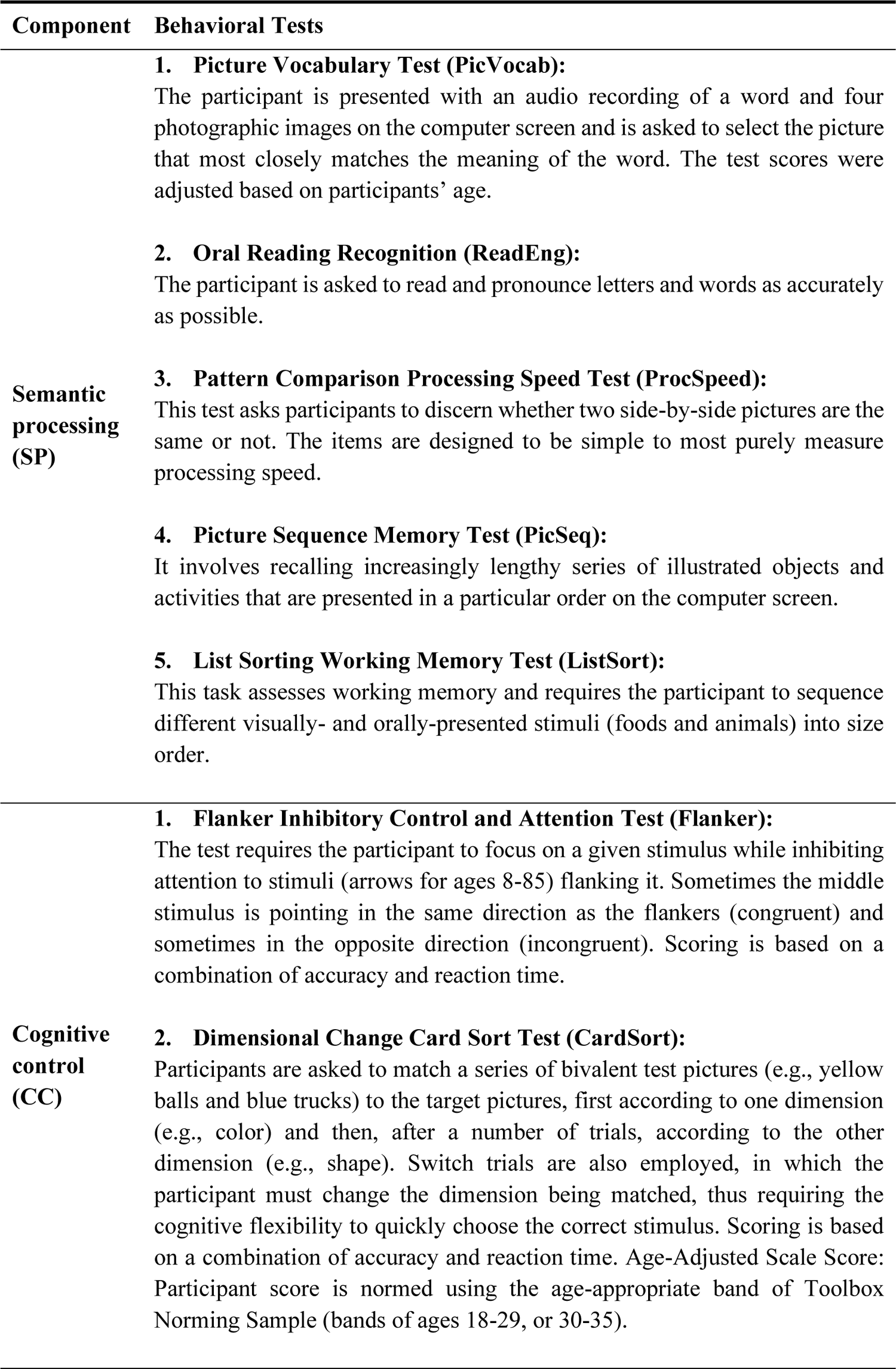
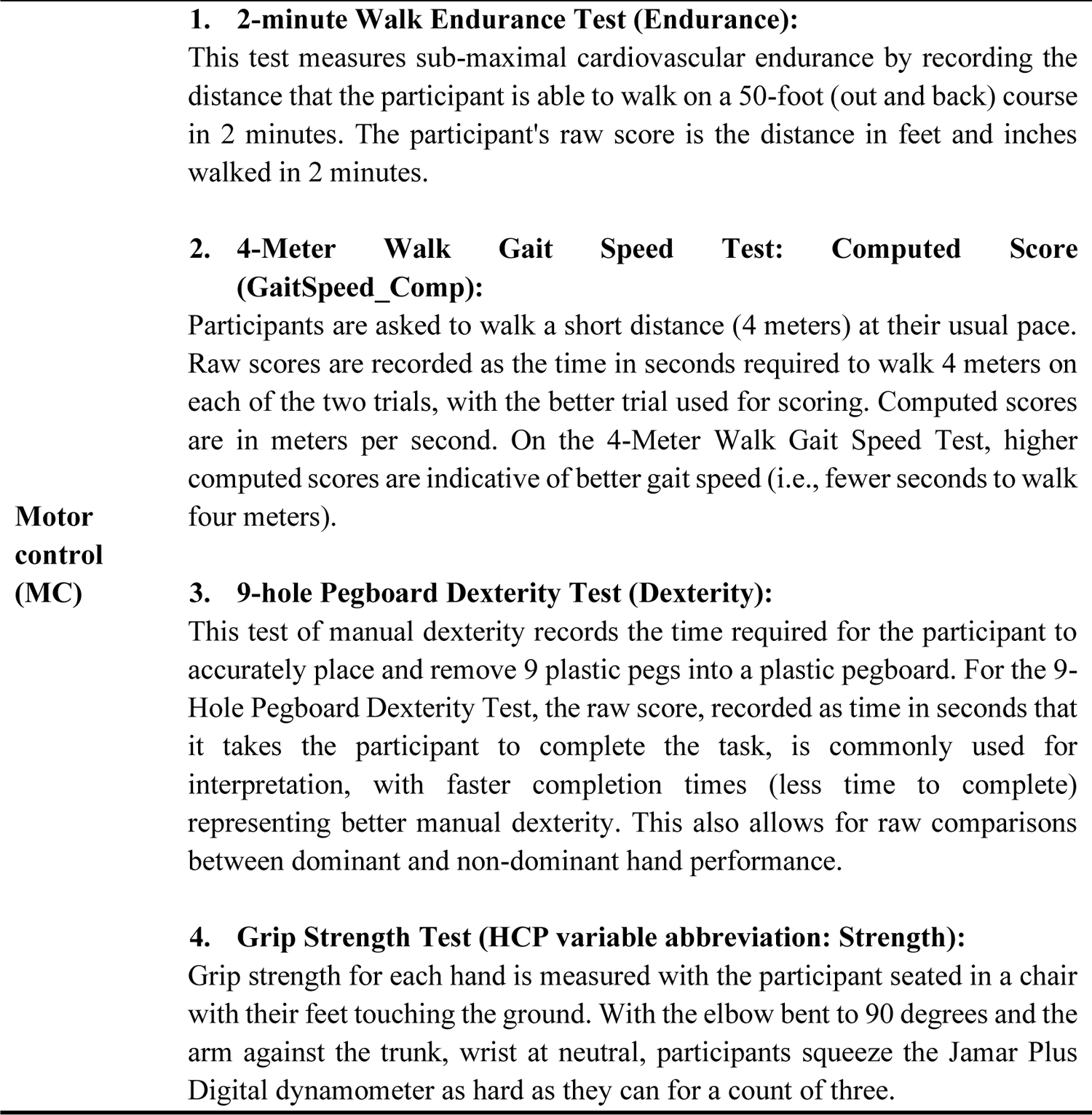
HCP behavioral assessments were selected to estimate three latent components with the confirmation factor analysis.

**Table S2.** Semantic brain template with ROI coordinates and region labels for the functional connectivity analysis.

| No. | x | y | z | ROI label | Module | Connector | Hub |
| --- | --- | --- | --- | --- | --- | --- | --- |
| 1 | -45 | -69 | 27 | Angular_L | DMN | DMN&PSN | – |
| 2 | -48 | -57 | 30 | Angular_L | PSN | DMN&PSN | – |
| 3 | -48 | -63 | 18 | Temporal_Mid_L | PSN | – | – |
| 4 | -3 | -57 | 18 | Precuneus_L | DMN | – | √ |
| 5 | -60 | -42 | -3 | Temporal_Mid_L | PSN | PSN&FPN | – |
| 6 | -45 | 30 | -9 | Frontal_Inf_Orb_L | PSN | – | √ |
| 7 | -60 | -51 | -9 | Temporal_Mid_L | FPN | – | – |
| 8 | -33 | -72 | 45 | Parietal_Inf_L | DMN | DMN&FPN | – |
| 9 | -36 | -78 | 33 | Occipital_Mid_L | DMN | – | √ |
| 10 | -27 | 30 | 45 | Frontal_Mid_L | DMN | – | – |
| 11 | -18 | 39 | 45 | Frontal_Sup_L | DMN | DMN&PSN | – |
| 12 | -54 | -63 | 9 | Temporal_Mid_L | PSN | – | – |
| 13 | 3 | -57 | 27 | Precuneus_R | DMN | – | √ |
| 14 | -42 | -69 | 39 | Angular_L | DMN | DMN&PSN&FPN | – |
| 15 | -30 | -36 | -15 | Fusiform_L | DMN | – | – |
| 16 | -57 | -48 | 6 | Temporal_Mid_L | PSN | – | – |
| 17 | 0 | -57 | 45 | Precuneus_L | DMN | – | – |
| 18 | 48 | -69 | 33 | Angular_R | DMN | – | √ |
| 19 | 57 | -57 | 24 | Temporal_Sup_R | DMN | DMN&PSN | – |
| 20 | -6 | -51 | 33 | Cingulum_Post_L | DMN | – | √ |
| 21 | -39 | -48 | 48 | Parietal_Inf_L | FPN | – | √ |
| 22 | -57 | -51 | 30 | SupraMarginal_L | PSN | – | √ |
| 23 | -39 | 24 | -18 | Frontal_Inf_Orb_L | PSN | – | – |
| 24 | -57 | -12 | -12 | Temporal_Mid_L | PSN | DMN&PSN | – |
| 25 | -6 | 48 | -6 | Frontal_Med_Orb_L | DMN | – | – |
| 26 | -57 | -54 | 15 | Temporal_Mid_L | PSN | – | – |
| 27 | -48 | 18 | 6 | Frontal_Inf_Tri_L | PSN | – | – |
| 28 | -12 | -60 | 12 | Calcarine_L | DMN | – | – |
| 29 | -45 | 27 | 21 | Frontal_Inf_Tri_L | FPN | – | – |
| 30 | -54 | -33 | -3 | Temporal_Mid_L | PSN | – | √ |
| 31 | -54 | -39 | 24 | Temporal_Sup_L | PSN | – | – |
| 32 | -48 | 9 | -21 | Temporal_Pole_Sup_L | PSN | – | – |
| 33 | -3 | -39 | 39 | Cingulum_Mid_L | DMN | – | √ |
| 34 | 63 | -39 | -3 | Temporal_Mid_R | PSN | – | – |
| 35 | -36 | 24 | 42 | Frontal_Mid_L | PSN | – | – |
| 36 | 54 | -48 | 12 | Temporal_Mid_R | PSN | – | – |
| 37 | 6 | 51 | -6 | Frontal_Med_Orb_R | DMN | – | – |
| 38 | -36 | 18 | 0 | Insula_L | PSN | – | – |
| 39 | -39 | 42 | -15 | Frontal_Inf_Orb_L | FPN | – | – |
| 40 | -54 | -3 | -24 | Temporal_Mid_L | PSN | DMN&PSN | – |
| 41 | -36 | -60 | 45 | Parietal_Inf_L | FPN | – | √ |
| 42 | -54 | -24 | -9 | Temporal_Mid_L | PSN | – | √ |
| 43 | 15 | -57 | 6 | Lingual_R | DMN | – | – |
| 44 | -54 | -45 | 39 | Parietal_Inf_L | FPN | – | – |
| 45 | -3 | -69 | 24 | Calcarine_L | DMN | – | √ |
| 46 | -18 | 30 | 51 | Frontal_Sup_L | DMN | DMN&PSN | – |
| 47 | 9 | -48 | 36 | Cingulum_Mid_R | DMN | – | √ |
| 48 | -3 | 48 | 21 | Frontal_Sup_Medial_L | PSN | – | – |
| 49 | -24 | 21 | 51 | Frontal_Mid_L | DMN | – | – |
| 50 | 54 | -63 | 15 | Temporal_Mid_R | DMN | – | – |
| 51 | 45 | -63 | 42 | Angular_R | DMN | – | – |
| 52 | -48 | -54 | 42 | Parietal_Inf_L | FPN | – | – |
| 53 | -24 | -78 | 42 | Occipital_Sup_L | DMN | – | – |
| 54 | -42 | -78 | 21 | Occipital_Mid_L | DMN | – | – |
| 55 | -51 | 18 | 24 | Frontal_Inf_Tri_L | FPN | – | √ |
| 56 | -42 | 15 | 45 | Frontal_Mid_L | PSN | DMN&PSN&FPN | – |
| 57 | -48 | 3 | -33 | Temporal_Inf_L | PSN | – | – |
| 58 | -36 | -54 | 57 | Parietal_Inf_L | FPN | – | – |
| 59 | 57 | -48 | 30 | SupraMarginal_R | PSN | – | – |
| 60 | 48 | -72 | 21 | Temporal_Mid_R | DMN | – | – |

Note. These coordinates, module, connector, and hub information are derived from Xu et al. 2016 (L = left, R = right, Mid = middle, Med = medial, Sup = superior, Inf = inferior, Orb = orbital, Tri = triangular).

**Table S3.** Significant selected edges (*P*-value = permutation-based FDR-corrected *P*) during the predictive modeling.

| No. | Type | Node 1 | Node 2 | Edge | $r$ | $P$-value |
| --- | --- | --- | --- | --- | --- | --- |
| 1 | positive | Temporal_Mid_L | Temporal_Mid_L | PSN-PSN | 0.220 | $1.24 \times 10^{-4}$ |
| 2 | positive | Frontal_Inf_Orb_L | Angular_L | PSN-FPN | 0.217 | $1.24 \times 10^{-4}$ |
| 3 | positive | Frontal_Inf_Orb_L | Temporal_Mid_L | PSN-FPN | 0.216 | $1.24 \times 10^{-4}$ |
| 4 | positive | Frontal_Inf_Orb_L | Temporal_Mid_L | PSN-PSN | 0.210 | $1.94 \times 10^{-4}$ |
| 5 | positive | Temporal_Mid_L | Temporal_Mid_L | PSN-PSN | 0.201 | $2.76 \times 10^{-4}$ |
| 6 | positive | Temporal_Pole_Sup_L | Temporal_Mid_L | PSN-PSN | 0.195 | $2.82 \times 10^{-4}$ |
| 7 | positive | Temporal_Mid_L | Temporal_Mid_L | PSN-PSN | 0.194 | $3.72 \times 10^{-4}$ |
| 8 | positive | Temporal_Mid_L | Cingulum_Post_L | PSN-DMN | 0.192 | $3.90 \times 10^{-4}$ |
| 9 | positive | Frontal_Inf_Orb_L | Cingulum_Post_L | FPN-DMN | 0.186 | $4.06 \times 10^{-4}$ |
| 10 | positive | Temporal_Mid_L | Temporal_Mid_L | PSN-PSN | 0.186 | $4.68 \times 10^{-4}$ |
| 11 | positive | Temporal_Pole_Sup_L | Temporal_Mid_L | PSN-PSN | 0.185 | $4.68 \times 10^{-4}$ |
| 12 | positive | Temporal_Pole_Sup_L | Temporal_Mid_L | PSN-PSN | 0.182 | $4.68 \times 10^{-4}$ |
| 13 | positive | Temporal_Mid_L | Temporal_Mid_L | PSN-PSN | 0.182 | $4.68 \times 10^{-4}$ |
| 14 | positive | Frontal_Inf_Orb_L | Frontal_Inf_Orb_L | PSN-FPN | 0.182 | $5.69 \times 10^{-4}$ |
| 15 | positive | Temporal_Pole_Sup_L | Angular_L | PSN-DMN | 0.181 | $6.05 \times 10^{-4}$ |
| 16 | positive | Temporal_Mid_L | Angular_L | PSN-DMN | 0.179 | $6.05 \times 10^{-4}$ |
| 17 | positive | Frontal_Mid_L | Frontal_Inf_Orb_L | PSN-FPN | 0.178 | $6.56 \times 10^{-4}$ |
| 18 | positive | Temporal_Mid_L | Frontal_Inf_Orb_L | PSN-PSN | 0.177 | $6.57 \times 10^{-4}$ |
| 19 | positive | Temporal_Mid_L | Frontal_Inf_Orb_L | PSN-PSN | 0.174 | $6.56 \times 10^{-4}$ |
| 20 | positive | Temporal_Mid_L | Temporal_Mid_L | PSN-PSN | 0.174 | $6.57 \times 10^{-4}$ |
| 21 | positive | Frontal_Inf_Orb_L | Temporal_Mid_L | PSN-PSN | 0.173 | $6.57 \times 10^{-4}$ |
| 22 | positive | Temporal_Mid_L | Temporal_Mid_L | PSN-PSN | 0.173 | $6.56 \times 10^{-4}$ |
| 23 | positive | Temporal_Pole_Sup_L | Temporal_Mid_L | PSN-PSN | 0.173 | $6.57 \times 10^{-4}$ |
| 24 | positive | Temporal_Mid_L | Precuneus_R | PSN-DMN | 0.173 | $6.57 \times 10^{-4}$ |
| 25 | positive | Temporal_Mid_L | Angular_L | PSN-DMN | 0.171 | $6.56 \times 10^{-4}$ |
| 26 | positive | Frontal_Inf_Orb_L | Temporal_Mid_L | PSN-FPN | 0.170 | $6.61 \times 10^{-4}$ |
| 27 | positive | Frontal_Inf_Orb_L | Temporal_Mid_L | PSN-PSN | 0.169 | $6.71 \times 10^{-4}$ |
| 28 | positive | Temporal_Mid_L | Frontal_Inf_Orb_L | PSN-PSN | 0.167 | $6.71 \times 10^{-4}$ |
| 29 | positive | Temporal_Pole_Sup_L | Frontal_Inf_Orb_L | PSN-PSN | 0.166 | $6.88 \times 10^{-4}$ |
| 30 | positive | Frontal_Inf_Orb_L | Temporal_Pole_Sup_L | PSN-FPN | 0.166 | $7.51 \times 10^{-4}$ |
| 31 | positive | Frontal_Inf_Orb_L | Temporal_Mid_L | PSN-FPN | 0.165 | $7.64 \times 10^{-4}$ |
| 32 | positive | Temporal_Mid_R | Temporal_Mid_L | PSN-DMN | 0.165 | $7.98 \times 10^{-4}$ |
| 33 | positive | Frontal_Inf_Orb_L | Cingulum_Post_L | PSN-DMN | 0.164 | $7.78 \times 10^{-4}$ |
| 34 | positive | Temporal_Mid_L | Angular_L | PSN-DMN | 0.162 | $8.22 \times 10^{-4}$ |
| 35 | positive | Frontal_Inf_Orb_L | Frontal_Inf_Orb_L | PSN-FPN | 0.162 | $8.32 \times 10^{-4}$ |
| 36 | positive | Temporal_Mid_R | Precuneus_R | DMN-DMN | 0.162 | $8.32 \times 10^{-4}$ |
| 37 | positive | Temporal_Pole_Sup_L | Temporal_Sup_R | PSN-DMN | 0.162 | $8.76 \times 10^{-4}$ |
| 38 | positive | Frontal_Inf_Orb_L | Angular_L | PSN-DMN | 0.160 | $9.12 \times 10^{-4}$ |
| 39 | positive | Frontal_Inf_Orb_L | Temporal_Mid_L | PSN-PSN | 0.159 | 9.39×10<sup>-4</sup> |
| 40 | positive | Temporal_Inf_L | Frontal_Inf_Orb_L | PSN-FPN | 0.156 | 9.93×10<sup>-4</sup> |
| 41 | negative | Temporal_Mid_L | Parietal_Inf_L | PSN-FPN | -0.223 | 1.24×10<sup>-4</sup> |
| 42 | negative | Temporal_Mid_L | Parietal_Inf_L | PSN-FPN | -0.205 | 1.94×10<sup>-4</sup> |
| 43 | negative | Parietal_Inf_L | Precuneus_R | FPN-DMN | -0.203 | 1.94×10<sup>-4</sup> |
| 44 | negative | Frontal_Inf_Orb_L | Parietal_Inf_L | FPN-FPN | -0.202 | 1.94×10<sup>-4</sup> |
| 45 | negative | Parietal_Inf_L | Temporal_Mid_L | PSN-FPN | -0.195 | 2.82×10<sup>-4</sup> |
| 46 | negative | Occipital_Mid_L | Angular_L | PSN-DMN | -0.190 | 3.72×10<sup>-4</sup> |
| 47 | negative | Parietal_Inf_L | Cingulum_Post_L | FPN-DMN | -0.189 | 3.72×10<sup>-4</sup> |
| 48 | negative | Occipital_Sup_L | Cingulum_Post_L | DMN-DMN | -0.173 | 6.56×10<sup>-4</sup> |
| 49 | negative | Parietal_Inf_L | Temporal_Sup_R | FPN-DMN | -0.171 | 6.71×10<sup>-4</sup> |
| 50 | negative | Parietal_Inf_L | Angular_L | FPN-DMN | -0.170 | 6.71×10<sup>-4</sup> |
| 51 | negative | Parietal_Inf_L | Temporal_Mid_L | PSN-FPN | -0.169 | 6.71×10<sup>-4</sup> |
| 52 | negative | Occipital_Mid_L | Precuneus_L | DMN-DMN | -0.169 | 6.71×10<sup>-4</sup> |
| 53 | negative | Parietal_Inf_L | Precuneus_R | FPN-DMN | -0.168 | 6.71×10<sup>-4</sup> |
| 54 | negative | Frontal_Inf_Orb_L | Parietal_Inf_L | PSN-FPN | -0.168 | 6.71×10<sup>-4</sup> |
| 55 | negative | Occipital_Sup_L | Precuneus_L | DMN-DMN | -0.166 | 6.88×10<sup>-4</sup> |
| 56 | negative | Parietal_Inf_L | Temporal_Mid_L | PSN-FPN | -0.164 | 7.98×10<sup>-4</sup> |
| 57 | negative | Calcarine_L | Temporal_Mid_L | PSN-DMN | -0.161 | 8.89×10<sup>-4</sup> |
| 58 | negative | Occipital_Mid_L | Frontal_Inf_Orb_L | PSN-DMN | -0.159 | 9.39×10<sup>-4</sup> |
| 59 | negative | Parietal_Inf_L | Precuneus_L | FPN-DMN | -0.158 | 9.93×10<sup>-4</sup> |
| 60 | negative | Occipital_Sup_L | Angular_L | PSN-DMN | -0.158 | 9.46×10<sup>-4</sup> |
| 61 | negative | Parietal_Inf_L | Temporal_Mid_L | PSN-FPN | -0.158 | 9.93×10<sup>-4</sup> |
| 62 | negative | Precuneus_R | Occipital_Mid_L | DMN-DMN | -0.157 | 9.93×10<sup>-4</sup> |

## Notes

### Competing Interest Statement

The authors have declared no competing interest.

